# SID-1 domains important for dsRNA import in *C. elegans*

**DOI:** 10.1101/156703

**Authors:** Jennifer S. Whangbo, Alexandra S. Weisman, Jiajie Lu, Jeiwook Chae, Craig P. Hunter

## Abstract

In the nematode *Caenorhabditis elegans,* RNA interference (RNAi) triggered by double-stranded RNA (dsRNA) spreads systemically to cause gene silencing throughout the organism and its progeny. We confirm that *Caenorhabditis* nematode SID-1 orthologs have dsRNA transport activity and demonstrate that the SID-1 paralog CHUP-1 does not transport dsRNA. Sequence comparison of these similar proteins, in conjunction with analysis of loss-of-function missense alleles identifies several conserved 2-7 amino acid microdomains within the extracellular domain that are important for dsRNA transport. Among these missense alleles, we identify and characterize a *sid-1* allele, *qt95*, which causes tissue-specific silencing defects most easily explained as a systemic RNAi export defect. However, we conclude from genetic and biochemical analyses that *sid-1(qt95)* disrupts only import and speculate that the apparent export defect is caused by the cumulative effect of sequentially impaired dsRNA import steps.Thus, consistent with previous studies, we fail to detect a requirement for *sid-1* in dsRNA export, but demonstrate for the first time that SID-1 functions in the intestine to support environmental RNAi.

## Introduction

In *C. elegans,* experimentally introduced double-stranded (ds)RNA triggers gene-specific silencing that spreads to silence the targeted gene in cells throughout the animal and in its progeny (Fire, Xu et al. 1998). The spread of gene silencing mediated by mobile intercellular silencing signals is termed systemic RNA interference (RNAi). Systemic RNAi has been described in plants and has been inferred in an abundance of other invertebrates (Jose and Hunter 2007). Furthermore, in *C. elegans* and some other animals, systemic gene silencing can be triggered by ingested dsRNA, a process known as environmental RNAi (eRNAi) (Whangbo and Hunter 2008). Genetic screens in *C. elegans* have identified **s**ystemic RNA**i d**efective (Sid) mutants (Winston, Molodowitch et al. 2002). The SID proteins identified by these mutations include two dsRNA transporters: SID-1 and SID-2 (Winston, Molodowitch et al. 2002, Feinberg and Hunter 2003, Winston, Sutherlin et al. 2007, McEwan, Weisman et al. 2012). SID-2 is a transmembrane protein expressed in the intestine and localized to the apical (lumenal) membrane (Winston *et al.* 2007) where it functions as an endocytic receptor for ingested dsRNA(McEwan, Weisman et al. 2012) As expected from this restricted expression pattern, *sid-2* mutant animals are defective for eRNAi, but are fully capable of initiating systemic RNAi in response to expressed or injected dsRNA (Winston *et al.* 2007). In contrast, SID-1 is a multi-pass transmembrane protein expressed in all cells sensitive to systemic RNAi (Winston, Molodowitch et al. 2002). SID-1 functions as a dsRNA channel to transport dsRNA across cellular membranes (Feinberg and Hunter 2003). Consistent with this broad expression, neither systemic nor eRNAi is detected in *sid-1* mutants (Winston, Sutherlin et al. 2007), although RNAi is robust within cells that express dsRNA or that are injected with dsRNA. This demonstrates that *sid-1* mutants are not defective for RNAi, but are only defective for the intercellular spread of silencing signals.

Analysis of *sid-1* genetic mosaic animals indicates that *sid-1* is required in the receiving cell, implicating a role for SID-1 in dsRNA import. Consistent with this finding, heterologous expression of *C. elegans* SID-1 in *Drosophila* S2 cells suggests that SID-1 is a dsRNA channel that passively transports extracellular dsRNA into cells (Feinberg and Hunter 2003). *C. elegans* experiments using tissue-specific SID-1 expression have shown that SID-1 is not required for export of silencing signals from cells expressing dsRNA (Jose, Smith et al. 2009). Although a role for SID-1 in dsRNA export has not been excluded, these experiments imply the existence of a SID-1-independent dsRNA export pathway.

Here, we report on a comprehensive functional and molecular analysis of all extant *sid-1* alleles recovered in our genetic screen. The recovery rate of duplicate single-nucleotide mutations in the coding region suggests that approximately 70% of phenotypically recoverable alleles have been discovered. All missense alleles alter amino acids conserved among inferred functional nematode SID-1 homologs, and all partial loss-of-function alleles are located within the N-terminal extracellular domain (ECD). Interestingly, all the ECD partial loss-of-function alleles are located within SID-1-specific 2-7 amino acids microdomains. One repeatedly recovered ECD missense substitution appears to selectively compromise dsRNA export from the intestine resulting in robust intestine-only eRNAi silencing. However, our focused analysis of this allele as well as more comprehensive genetic mosaic analyses suggest that this apparent export defect likely represents two or more successive compromised dsRNA import steps. These analyses provide evidence that amino acids important for efficient dsRNA transport are conserved among SID-1 homologs in nematodes and that SID-1-dependent dsRNA import into the intestine is a key step for efficient dsRNA delivery to other tissues providing insight into the mechanisms of intercellular dsRNA transport. This analysis resolves the role of *sid-1* in dsRNA export, provides evidence that sid-1 functions during eRNA in the intestine, and identifies specific SID-1 domains important for dsRNA transport.

## Results

### C. *brenneri* SID-1 is a functional dsRNA transporter while the SID-1 homolog TAG-130/CHUP-1 does not transport dsRNA

SID-1 is highly conserved among the five sequenced *Caenorhabditis* nematodes, yet *C. brenneri* lacks systemic RNAi in response to injected and ingested dsRNA (Winston, Sutherlin et al. 2007). This is particularly curious as *C. brenneri* SID-1 (CBN-SID-1) is most similar to *C. elegans* SID-1 (CEL-SID-1) in pairwise comparisons of the *Caenorhabditis* homologs (Figure S1, Figure 1B). Although well conserved, CBN-SID-1may be non-functional for dsRNA transport, inadequately expressed, or SID-1-independent activities required for systemic RNAi may be lacking in *C. brenneri*. To address these possibilities, we first asked whether *Cbn-sid-1* could complement a *C. elegans sid-1* mutant to restore systemic RNAi. We tested animals with a feeding RNAi assay using bacteria that express dsRNA targeting either *act-5* (intestine) or *unc-22* (muscle). Both of these genes are associated with an easily scored, fully penetrant RNAi knockdown phenotype in wild-type worms (see Materials and Methods). We found that a *C. brenneri sid-1* genomic fragment could rescue the null Sid phenotype of *sid-1(qt9) C. elegans* mutants. Specifically, the co-injection marker positive animals showed 100% sensitivity to *unc-22* eRNAi (twitching phenotype) and *act-5* eRNAi (larval arrest phenotype), while the co-injection marker negative animals showed 100% resistance (Figure 1C). Thus, in the context of *C. elegans*, the CBN-SID-1 protein enables sufficient dsRNA transport for detectable eRNAi. To determine whether CBN-SID-1 transports dsRNA as efficiently as CEL-SID-1, we compared dsRNA uptake between *Drosophila S2* cells expressing CEL-SID-1 and CBN-SID-1 (Figure 1D). Consistent with the high sequence conservation, CBN-SID-1 functions as a robust dsRNA transporter in this *in vitro* assay. Thus, we conclude that CBN-SID-1 is a functional SID-1 ortholog, and that a defect in dsRNA transport activity cannot explain the lack of systemic RNAi in *C. brenneri.* Therefore, all five sequenced SID-1 homologs are functional for systemic RNAi and may be classified as orthologs (Winston et al., 2007; Figure 1; pers. comm. CPH for C. japonica).

**Figure 1.**
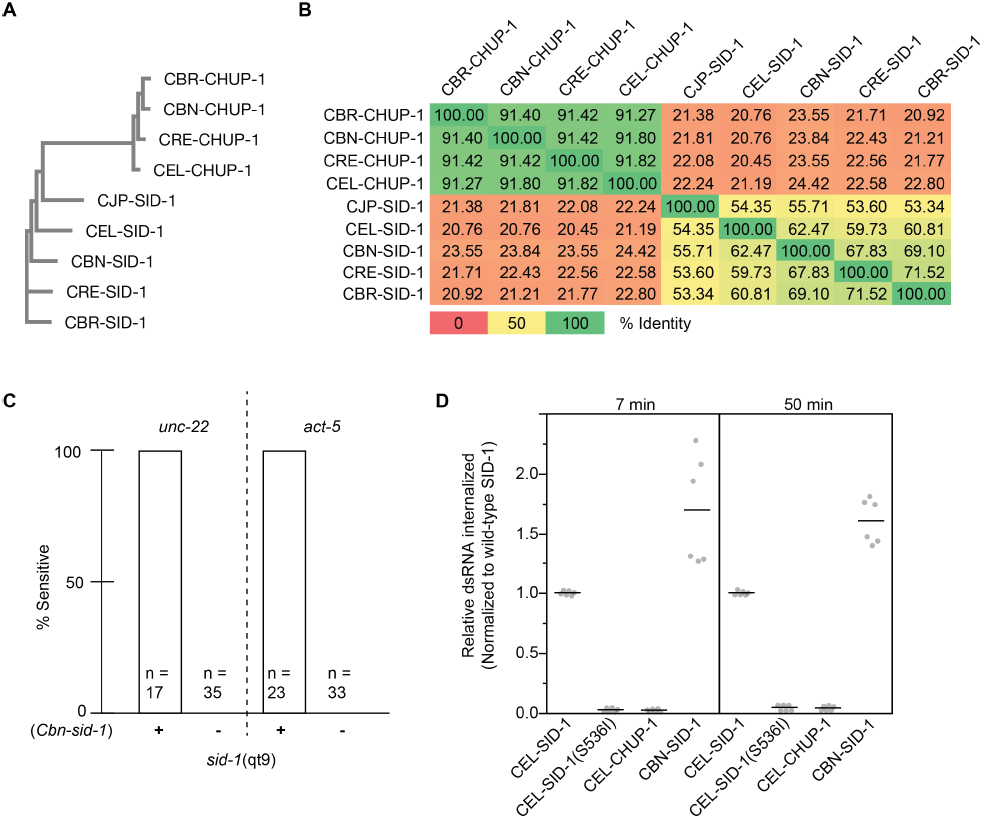
Sequence and functional relationships among SID-1 homologs. (A) Phylogenetic tree of nine Caenorhabditis SID-1 homologs (MUSCLE v3.8). (B) Pairwise percent identity heat matrix of homologs. (C) *sid-1(qt9)* animals with or without an extrachromosomal array containing a *cbn-sid-1* genomic fragment were tested for RNAi sensitivity on *unc-22* or *act-5* RNAi food. (D) Internalization of ^32^P-labeled 500 bp dsRNA into Drosophila S2 cells expressing the indicated SID-1 constructs. Line represents mean value of 2 biological replicates.

All sequenced *Caenorhabditis* species contain additional SID-1 homologs, including the highly conserved gene *tag-130/chup-1* (Tomoyasu, Miller et al. 2008), (Figure 1A). The inability of *tag-130/chup-1* to complement *sid-1* mutants may reflect its more limited expression or its more divergent sequence. The four full-length *Caenorhabditis* CHUP-1 homologs share greater than 90% sequence identity with each other (pairwise comparison), but share less than 25 % identity between any SID-1 and any CHUP-1 homolog (Figure 1B). Consistent with the apparent lack of a role in systemic RNAi, the *C. elegans* CHUP-1 homolog (CEL-CHUP-1) shows no detectable RNA transport activity when expressed in *Drosophila* S2 cells (Figure 1D). Therefore, we conclude that *sid-1* and *tag-130/chup-1* are paralogs with distinct activities.

### Missense mutations map to the extracellular and transmembrane domains

Despite the paucity of aligned identical amino acids among the nine homologs, the predicted CHUP-1 topology matches the partially determined and predicted SID-1 topology quite well, thus we presume that amino acids that are conserved between the two proteins are likely structurally important, while amino acids that are conserved only among the SID-1 orthologs are likely functionally important for dsRNA transport (Figure S1, Figure 2D). To determine which amino acids and protein domains are important for dsRNA transport we sequenced all independently isolated extant *sid-1* alleles from our original Sid screen, which together with previously described alleles, identified 49 point mutations and seven small deletions at the *sid-1* locus (Table S1, Figure 2AC) (Winston et al., 2002). In conjunction with the sequencing effort we tested these mutants via environmental RNAi (eRNAi) on a panel of foods targeting five distinct tissues (Table S2, Figure 3). The Sid phenotype of an isolate was defined as “strong” if fewer than10% of animals were sensitive to each tested trigger, and all others were defined as “weak”. All alleles also tested as recessive (data not shown).

**Figure 2.**
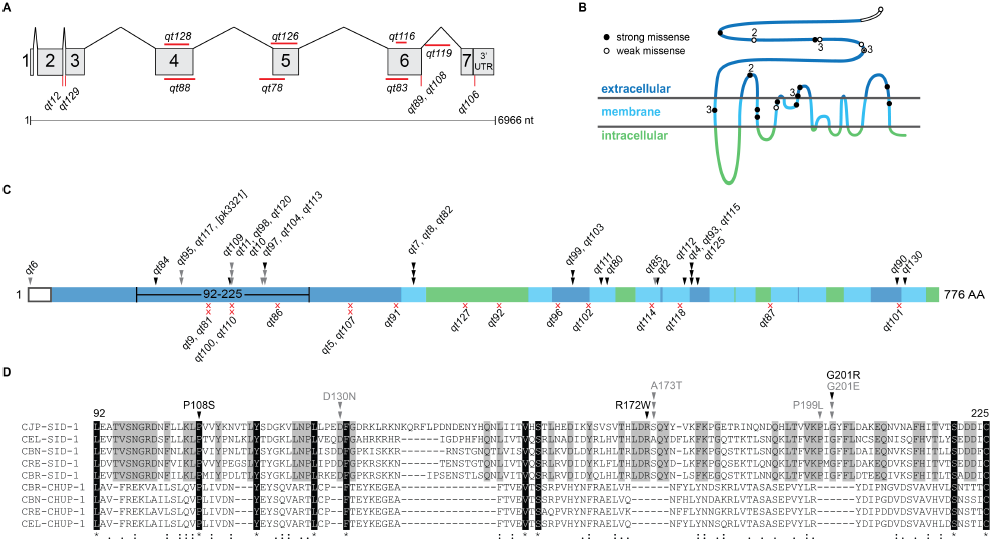
Location and nature of recovered *sid-1* alleles. (A) Genic representation of recovered non-coding and deletion *sid-1* alleles. (B) SID-1 topological model annotated with strong (black) and weak (white) missense alleles. Numbers indicate multiple independent isolates. (C) Distribution and number of independently recovered strong (black) and weak (grey) missense (arrowheads) and nonsense (red x) alleles. Topological domains are shaded as in panel B. (D) Section of a multiple sequence alignment (Fig S1) corresponding to the region of the extracellular N-terminus (amino acids 92-225) with frequently recovered missense alleles. Residues identically conserved in all nine aligned sequences are highlighted in black; residues identically conserved only in the five putative SID-1 orthologs are highlighted in grey.Similarity of the nine homologs is indicated on the bottom line. Arrowheads are as described in panel C.

**Figure 3.**
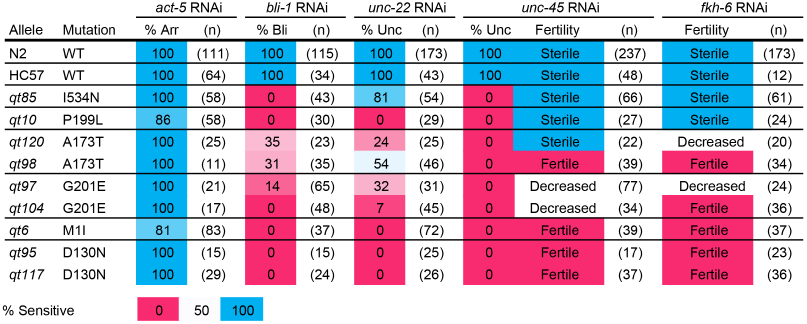
Weak *sid-1* alleles on tissue-specific environmental RNAi. N2, wild type strain. HC57, background strain for recovery of *sid-1* mutants. Bli, blistered cuticle; Unc, uncoordinated (*unc-45* RNAi animals are paralyzed, *unc-22* RNAi animals are twitching). Fertility was scored qualitatively: sterile (no progeny), fertile (many progeny), decreased (few progeny). (n), number of animals scored. ND, not determined.

We recovered 16 nonsense alleles at 13 sites (10 singles and 3 duplicate isolates) and 28 missense alleles at 19 positions (13 single, 3 double, and 3 triple isolates). This frequency of multiple isolates indicates that this genetic screen is about 70% saturated for *sid-1* alleles with a readily detected Sid phenotype. More specifically, this fraction of duplicate recovered nonsense alleles (mean nonsense alleles/site = 1.23) leads to a Poisson estimate (P_0_1=e^-m^, where m = mean allele freq/site) of five to six additional recoverable nonsense alleles and similarly six to seven additional recoverable missense alleles (mean alleles/per site = 1.47) (Figure 2C). The nonsense alleles and the majority of the missense alleles had very strong loss-of-function Sid phenotypes.The nonsense alleles were distributed throughout the protein and in conjunction with the recovered small deletions we conclude that the *sid-1* null phenotype is fully viable and fertile.

In contrast to the nonsense alleles, missense alleles were recovered only in predicted extracellular or transmembrane domains. The missense alleles with strong Sid phenotypes target amino acids conserved in both SID-1 and CHUP-1 homologs; 10 of 13 strong missense allele sites alter amino acids identical (7) or conserved between SID-1 and CHUP-1. This observation is consistent with the inference that amino acids conserved between SID-1 and CHUP-1 homologs are important for structure/topology of the proteins. In contrast four of six missense allele sites with weak Sid phenotypes target amino acids within SID-1-specific stretches of amino acids, or microdomains, that that are not present in the CHUP-1 homologs (Figure S1, Figure 2D). These microdomains, which were either lost from CHUP-1 or gained in SID-1 homologs, are located within the ECD. Furthermore, despite the weak Sid phenotype, we recovered multiple independent alleles within each microdomain (Figure 2D, Table S1). This mutational and conservation data strongly implicates these domains in dsRNA-specific functions.

The partial loss-of-function alleles, particularly since they cluster in the ECD, may define subtle roles for *sid-1* in dsRNA transport. Curiously, on the panel of feeding RNAi clones targeting genes expressed in a variety tissues, these weak alleles did not show a simple pattern of reduced sensitivity to systemic RNAi across all tissues (Figure 3).Instead, several alleles exhibit tissue-specific deficiencies in response to eRNAi. For example, *qt10* and *qt85* are both sensitive to eRNAi silencing the intestinal gene *act-5*, and gonad/germline genes *fkh-6* and *unc-45*, and both resistant to silencing the cuticle gene *bli-5.* However, *qt10* animals are completely resistant to silencing the muscle gene *unc-22*, while *qt85* animals are sensitive. Two of the weak alleles *qt95* and *qt117*, which encode the same molecular lesion (D130N), exhibit a particularly interesting pattern of eRNAi sensitivity. Mutants with this lesion are sensitive to the intestinal *act-5* RNAi but fully resistant to all other RNAi triggers. A third independently isolated allele of *sid-1 (pk3321)* also shares this molecular lesion (Tijsterman, May et al. 2004).

Because the intestine is in direct contact with ingested dsRNA, this particular phenotype generates many new questions about the role of SID-1 in eRNAi. Uptake of ingested dsRNA requires both SID-1 and the intestine-restricted SID-2 for dsRNA transport from the gut lumen into the cytoplasm of intestinal cells (Winston, Sutherlin et al. 2007). Thus, this lesion may selectively disrupt *sid-1* function in non-intestinal cells,while retaining *sid-2*-dependent *sid-1* functions required for the uptake of ingested dsRNA. Alternatively, the intestine could be inherently more sensitive to eRNAi than other tissues or the intestine-specific *act-5* food may be less potent than the other tissue-specific dsRNA triggers. We selected *qt95* for all further studies and hoped to gain insight into the process of eRNAi through in depth characterization of this allele.

### The intestine is not more sensitive to eRNAi

To test the possibility that the intestine is inherently more sensitive to ingested dsRNA than other tissues we performed an experiment that would enable simultaneous visualization of RNAi-mediated silencing in multiple tissues. Wild-type animals carrying the *sur-5::gfp* transgene, which expresses GFP in all somatic nuclei, were placed on bacteria expressing *gfp* dsRNA as L4s and scored for GFP expression. If the intestine was more sensitive to RNAi we would expect that silencing of GFP in the intestine, but not other tissues, would occur earlier or at lower concentrations of *gfp* dsRNA-expressing bacteria. After 48 hours on the *gfp* trigger animals exhibited efficient silencing in both intestinal and hypodermal nuclei. Experiments scoring GFP expression at 24 hours were performed, but only uniform intermediate silencing (dimming) of intestinal and hypodermal nuclei was observed (Figure 4AB). To test whether lower concentrations would enable visualization of differential silencing, the RNAi assay was repeated as a dilution series using bacteria expressing L4440, empty vector, to dilute the *gfp* trigger. At dilutions that led to any silencing in any tissue, we found that silencing of GFP expression in the intestine occurred concurrently with silencing in other tissues (Figure 4C). Thus, in wild-type, proximal tissues (e.g. intestine) are not more sensitive to eRNAi than distal tissues (e.g. muscle).

**Figure 4.**
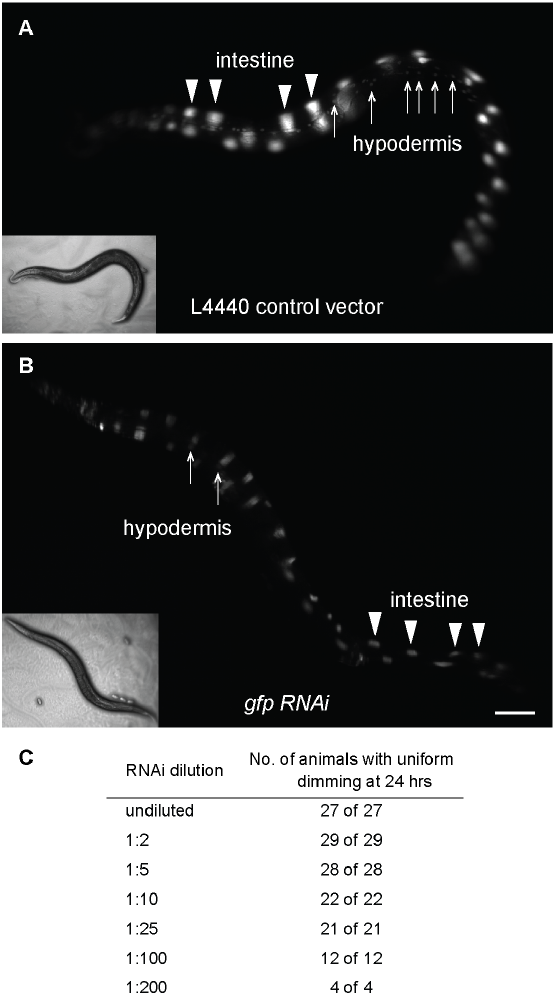
The intestine is not more sensitive to feeding RNAi than other tissues. Wild-type L4 animals were placed on a dilution series of feeding *gfp* RNAi and scored for silencing in the intestinal nuclei and hypodermal nuclei at 24 hours. (A) Wild-type animals at 24 hours on bacteria expressing the L4440 control RNAi vector. (B) Wild-type animals at 24 hours on bacteria expressing undiluted *gfp* RNAi.(C) Uniform silencing of intestinal and hypodermal nuclei in response to a dilution series of *gfp* RNAi in wild-type animals.

### *sid-1(qt95)* is functional for dsRNA import in all tissues

The *sid-1(qt95)* intestine-specific eRNAi sensitivity phenotype suggests that SID-1(D130N) is defective for dsRNA import in non-intestinal tissues. To determine whether *sid-1(qt95)* mutant animals are competent for import of dsRNA into tissues other than the intestine, we injected dsRNA directly into the pseudocoelom (PC). This fluid-filled body cavity separates the body wall of nematodes (the cuticle, hypodermis, excretory system, neurons and muscles) from the internal tissues (the pharynx, intestine and gonad). We once again utilized the *sur-5::gfp* transgene to visualize RNAi-mediated silencing in multiple tissues simultaneously. *gfp* dsRNA injected into the PC of animals wild-type for *sid-1* resulted in complete silencing of GFP expression in all tissues within 48 hours following injection (Figure 5A). In contrast, pseudocoelomic delivery of dsRNA into animals carrying a null allele, *sid-1(qt9),* did not lead to silencing in any tissue (Figure 5C). Injection of *gfp* dsRNA into the PC of *sid-1(qt95)* animals resulted in complete silencing of *gfp* in the entire animal in 15/22 animals (Figure 5E). The remaining 7/22 animals showed complete silencing of GFP expression in approximately half of the body, proximal to the injection site. This incomplete penetrance may indicate a partial dsRNA import defect. Similar results were obtained following pseudocoelomic injection of dsRNA homologous to an endogenous muscle-specific gene. Injecting *unc-22* dsRNA into the pseudocoelom of *sid-1(qt95)* animals produced characteristic Unc-22 twitching (Fire, Xu et al. 1998) in 7/7 injected animals and their progeny, whereas pseudocoelomic injection of *unc-22* dsRNA into *sid-1(qt9)* animals did not produce any twitching in the injected animals or their progeny. Thus, unlike *sid-1(qt9)* animals, which are resistant to injection of any dsRNA into the PC (Figure 5C; (Winston, Sutherlin et al. 2007)), *sid-1(qt95)* animals are able to import silencing signals from the PC into adjacent tissues.

**Figure 5.**
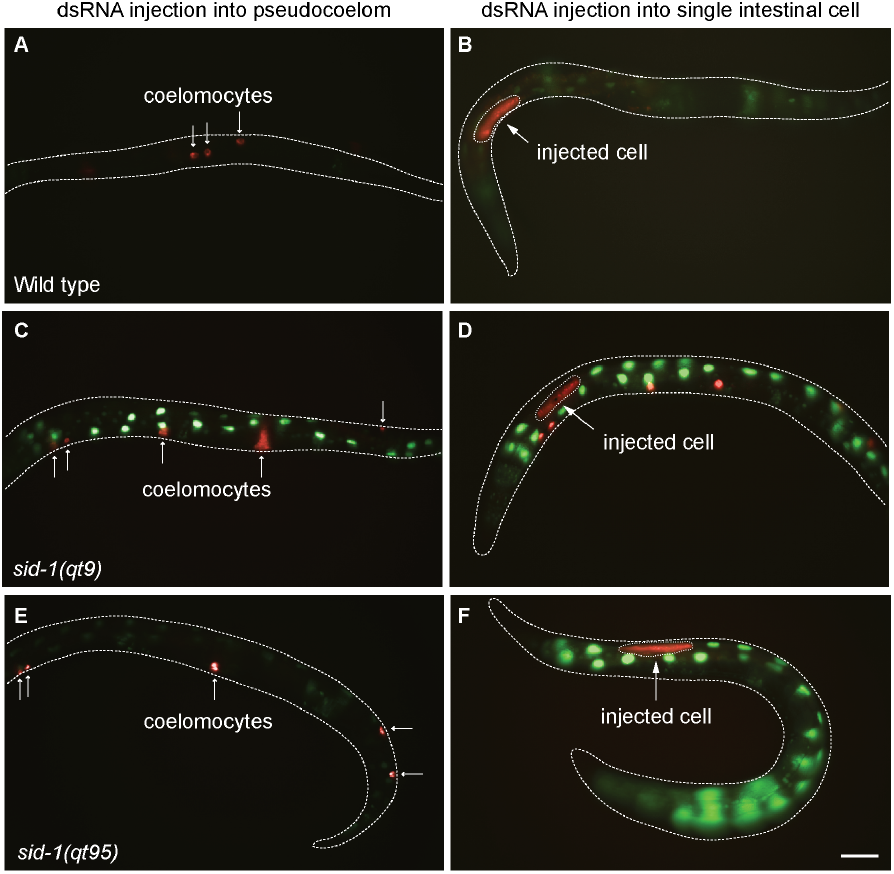
DsRNA import in *sid-1(qt95)* animals. Wild-type, *sid-1(qt9),* and *sid-1(qt95)* expressing sur-5::*gfp* were co-injected with *gfp* dsRNA and Alexa Fluor 555 Dextran (AD) in either the pseudocoelom (PC) or a single intestinal cell and imaged 48 hours later. In PC injected animals the AD (red) is only detected in coelomocytes (small arrows) (A, C, E). In intestinal cell injected animals that did not leak, the AD is restricted to the injected cell (larger arrow) (B, F). If the cell leaked, then AD is readily detected in coelomocytes as well (D). GFP expression was silenced in 20/20 PC injected WT animals (A). GFP expression was not silenced 9/9 PC injected sid-1(qt9) animals (C). GFP expression was silenced in 15/22 PC injected sid-1(qt95) E) and silenced in approximately half the animal in 7/22 PC injected animals. GFP expression was silenced in all intestinal cells in 5/11 single-intestinal cell injected WT animals (B) and in approximately half the intestinal cells in 6/11 animals. GFP was silenced only in the injected intestinal cell in 5/5 sid-1(qt9) animals (D) and 14/14 sid-1(qt95) animals (F). Injected intestinal cells in (B), (D) and (F) are outlined. Scale bar, 100 μm.

### *sid-1* is not required for dsRNA export from injected intestinal cells

Our observations to this point show that in *sid-1(qt95)* animals, eRNAi leads to effective gene silencing in the intestine, but not in other tissues; that bypassing the intestine with dsRNA injection into the pseudocoelom causes efficient silencing in all tested tissues. One explanation for these results is that *sid-1(qt95)* blocks export of silencing signals from the intestine. The initial genetic mosaic analysis of *sid-1* function, which demonstrated that *sid-1* is required for import of eRNAi triggers, did not address a role in the export of silencing signals (Winston, Molodowitch et al. 2002). A subsequent report showed that SID-1 is not required for export of pharyngeal-expressed dsRNA (Jose, Smith et al. 2009). However, it is possible that the requirement for *sid-1* for dsRNA export is specific to eRNAi.

Consistent with this possibility, injection of *gfp* dsRNA into single intestinal cells of *sid-1(qt95)* animals carrying the *sur-5::gfp* transgene led to silencing of *gfp* expression in only the injected cell in 9/11 animals (Figure 5F). In the remaining two animals, there was spread of silencing from the injected cell only to adjacent intestinal cells, but not outside of the intestine. This may indicate that the *qt95* mutation does not completely block export, or may be due to accidental puncturing of adjacent cells or leakage of dsRNA by cell rupture during dsRNA injection. As expected, injection into single intestinal cells of wild-type animals led to diffuse spread of silencing, whereas injection into single intestinal cells of *sid-1(qt9)* null mutants led to silencing only in the injected cell (Figure 5B, D).

Although these results support a role for *sid-1* in the export of injected dsRNA from the intestine, a more definitive test for a *sid-1* requirement in dsRNA export is dsRNA injection into *sid-1(null)* mosaic intestines. Specifically, we injected *gfp* dsRNA into *sid-1(qt9)*(null) intestinal cells and asked whether *gfp* expression in adjacent and nearby *sid-1(+)* cells was silenced. We generated *sid-1(qt9)* mosaic animals by spontaneous loss of an extrachromosomal array during embryonic cell divisions (Herman and Hedgecock 1990, Yochem, Gu et al. 1998). The extrachromosomal array was composed of a wild-type *sid-1* genomic DNA PCR product that rescues *sid-1* (Winston, Molodowitch et al. 2002) and *sur-5::GFP* plasmid. GFP positive animals from this line are sensitive to eRNAi, while GFP negative animals are resistant.

For the *gfp* dsRNA injection experiments, animals with a mixture of GFP positive and GFP negative intestinal cells were selected. To ensure that only properly injected animals were scored, we included Alexa Fluor 555 conjugated to dextran in the *gfp* dsRNA injection mix. Any dextran-Alexa Fluor 555 that leaks into the PC space is concentrated in coelomocytes identifying animals in which dsRNA was not injected solely into a single intestinal cell. In five independent lines we found that injecting *gfp* dsRNA into *sid-1(qt9)* non-GFP cells silenced GFP expression in all *sid-1(+)* intestinal cells (Figure 6). This result indicates that *sid-1* is not required for export of injected dsRNA, and is in agreement with the *sid-1*-independent export of expressed dsRNA (Jose, Smith et al. 2009). Thus, an explanation for the unusual *qt95* phenotype must be found elsewhere.

**Figure 6.**
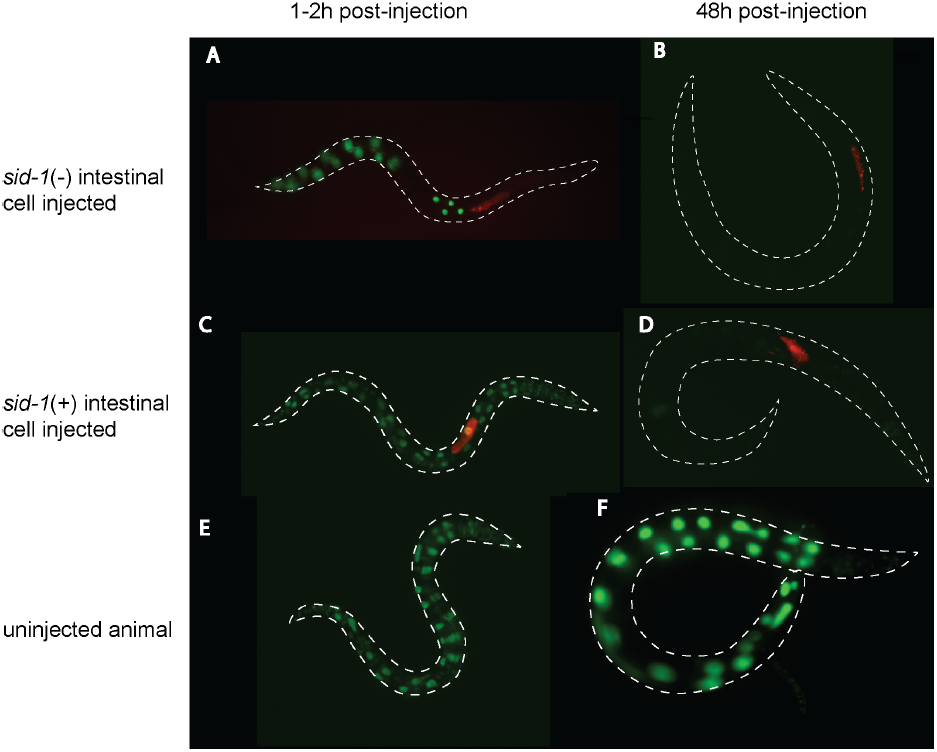
*sid-1* is not required for export from the intestine. Young adult wild-type or *sid-1* mosaic animals were co-injected with *gfp* dsRNA and Alexa Fluor 555 dye conjugated to 10,000 MW inert Dextran (AD), recovered at 20°C. The animals were photographed 1-2 hours (left) and 48 hours (right) post-injection. Merged green and red channel images show GFP expression and AD, respectively. (A,B) Intestinal *gfp* expression in *sid-1* mosaic animal injected with *gfp* dsRNA in *sid-1*(-) intestinal cell (A) before and (B) after silencing. Similar silencing was observed in multiple mosaics isolated from five independent lines. (C,D) Intestinal *gfp* expression in control animal injected in *sid-1*(+) intestinal cell (C) before and (D) after silencing. (E, F) Concurrently staged uninjected control animals. All *gfp* images are adjusted identically.

### *sid-1(qt95)* is sensitive to dsRNA dose

The *sid-1(qt95)* allele confers complete resistance to eRNAi in the muscle, hypodermis, somatic gonad and germline, yet these tissues are sensitive to systemic RNAi initiated by injection of dsRNA into the pseudocoelom. As described above, *gfp* dsRNA injections into the intestines of mosaic *sid-1(qt9)* animals have ruled out a requirement for *sid-1* in dsRNA export. Although consistent with a tissue-to-tissue transport defect, other explanations, singly or in combination may also explain the observed *sid-1(qt95)* phenotype.

Rather than differential tissue sensitivity, could the tissue-specific differences revealed by the weak *sid-1* mutants simply reflect gene-specific differences in RNAi sensitivity? A recent study compared the relative penetrance of RNAi sensitivity at varying doses of dsRNA-expressing bacteria and demonstrated that some ingested RNAi triggers are more potent than others (Zhuang and Hunter 2012). Thus, it is possible that the activity of *sid-1(qt95)* may be sufficient to enable silencing by a potent RNAi trigger, such as *act-5,* but not a less potent trigger, such as *fkh-6*.

To determine whether differences in gene-specific RNAi potency can explain the tissue-specific responses observed in *sid-1(qt95)* mutants, we repeated eRNAi assays using a single dsRNA trigger (*gfp* dsRNA) targeting the same gene expressed in multiple tissues. Animals carrying the *sur-5::gfp* transgene were placed on bacteria expressing *gfp* dsRNA as L4 larvae and then scored for *gfp* expression after 48 hours of feeding. Wild-type animals carrying the *sur-5::gfp* transgene were completely silenced in all tissues scored except for neurons, which are refractory to RNAi (Figure 7B). As expected for a null allele, *sid-1(qt9)* animals carrying the *sur-5::gfp* transgene did not show *gfp* silencing in any tissues after 48 hours (Figure 7D). Consistent with the previous observations, *sid-1(qt95)* animals carrying the *sur-5::gfp* transgene showed gfp silencing in the intestine, but not in other tissues (Figure 7F). Thus, differences in RNAi trigger efficacy do not readily explain the sid-1(qt95) phenotype.

**Figure 7.**
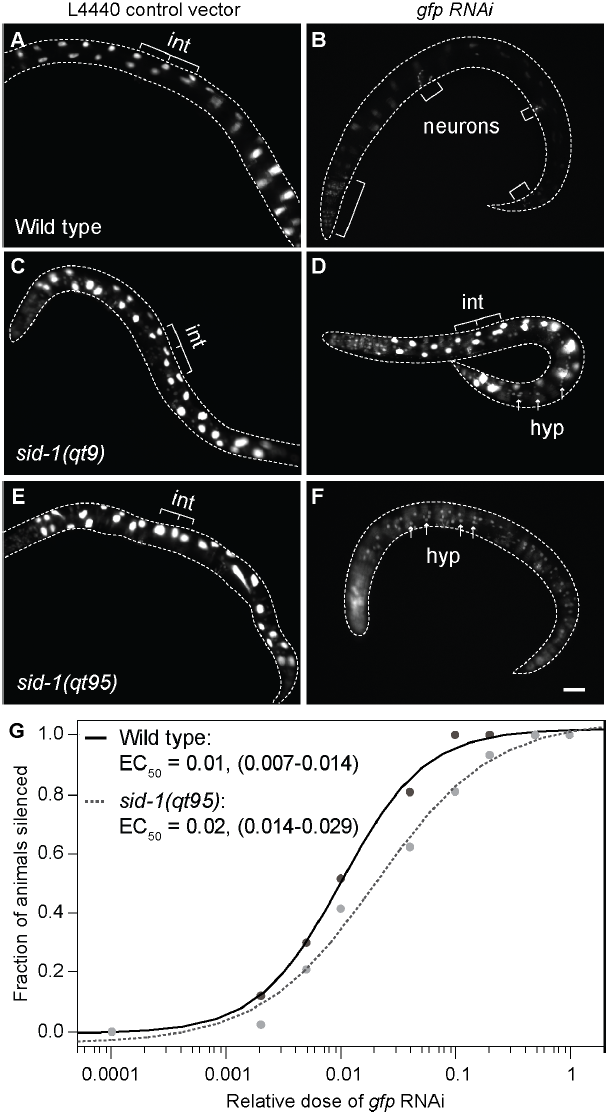
Tissue-specific *gfp* eRNAi silencing in *sid-1(qt95)* animals. *sur-5::gfp* L4 animals were placed on L4440 (control) vector or *gfp* RNAi bacteria at 20°C and scored at 48 hours for *gfp* expression. n > 25 animals for each condition. Scale bar, 100 μm. (A,B) Wild-type animals grown on *gfp* RNAi show silencing of gfp in all tissues except neurons, which are refractory to RNAi. (C, D) *sid-1(qt9)* animals are fully defective in environmental RNAi. (E, F) *sid-1(qt95)* animals show silencing of intestinal nuclei only.(G) *gfp* silencing in response to a dilution series of *gfp* RNAi bacteria in wild-type and *sid-1(qt95)* animals. The fraction of animals with intestinal *gfp* silencing at each dilution and the apparent half maximal effective concentration (EC50) with 95% confidence interval in parentheses is shown. n > 20 animals for each dilution. int, intestinal nuclei. hyp, hypodermal nuclei.

Finally, the suppression of the *sid-1(qt95)* defect by pseudocoelomic injected dsRNA may simply reflect the much higher dsRNA concentration or dose compared to ingested dsRNA. Similarly, the available dose of ingested dsRNA may be higher in the intestine than other tissues. To test the sensitivity of *sid-1(qt95)* worms to varying dsRNA doses, we compared the silencing efficiency of wild-type and *sid-1(qt95)* animals carrying the *sur-5::gfp* transgene in response to a dilution series of *gfp* eRNAi. The apparent half maximal effective concentration (EC_50_) for *gfp* silencing in the intestinal nuclei was a *gfp* RNAi dilution of 0.01 for wild-type animals and 0.02 for *sid-1(qt95)* animals (Figure 7G). Using the analysis of means (ANOM) for the EC_50_ estimates, the observed silencing wild-type and *sid-1(qt95)* animals was significantly different at an alpha level of 0.05, but not at an alpha level of 0.01. Thus, eRNAi silencing in the intestine may show a difference in dose sensitivity between *sid-1(qt95)* and wild type. How this dose sensitivity could lead to tissue-specific difference in silencing in response to full-strength food requires additional explanation.

### *Drosophila* S2 cells expressing SID-1(D130N) have reduced uptake activity

The partial silencing proximal to the PC injection site of *sid-1(qt95)* animals, and the subtle difference in dosage sensitivity of wild-type and *sid-1(qt95)* animals suggest that the *sid-1(qt95)* mutants may be partially impaired in dsRNA importing steps. Our *in vitro* dsRNA uptake assay is our most sensitive assay for SID-1 function. Therefore, we used this assay to directly test whether the product of the *sid-1(qt95)*, SID-1(D130N), retained dsRNA transport activity.

We first performed a time-course experiment, measuring import of radiolabeled dsRNA into *Drosophila* S2 cells transfected with WT or mutant SID-1 at regularly spaced time points in order to calculate dsRNA uptake rates. The results of this kinetic assay and our standard uptake assay indicated that SID-1(D130N)-expressing S2 cells performed at 5-10% of the rate of SID-1(WT)-expressing S2 cells (Figure 8A, B). This confirms that the dsRNA import activity of SID-1(D130N) channels is partially impaired. Consistent with a passive transport mechanism, both extending the incubation time and increasing the dsRNA concentration increased the amount of dsRNA transported into cells, but under both conditions SID-1(D130N) expressing cells were ten-fold less active than wild-type SID-1 expressing cells but more active than the non-functional SID-1(S536I) expressing cells (Figure 8B).

**Figure 8.**
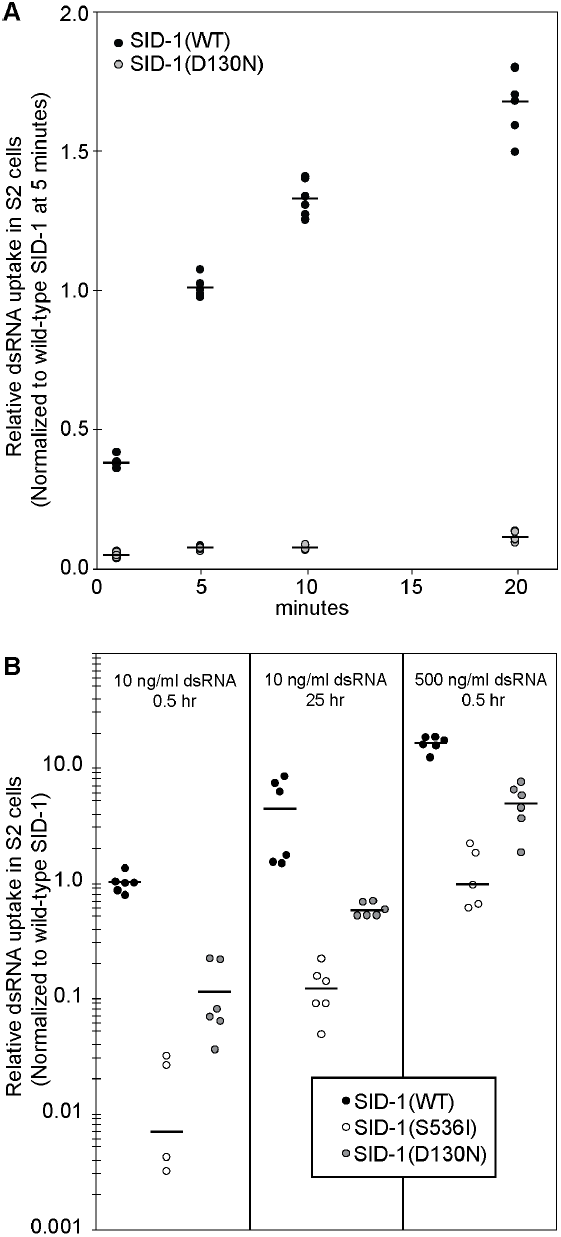
dsRNA import activity of SID-1(D130N). Internalization of ^32^P-labeled 500 bp dsRNA in Drosophila S2 cells expressing the indicated SID-1 construct. (A) Time course assays performed at room temperature. (B) For duration and concentration assays cells were pre-incubated at 4C for 0.5 hours, and assays were performed at 4°C. Line is the mean of two biological replicates and three technical replicates (Two S536I replicates were negative when normalized to background and are not plotted).

### Expression of WT *sid-1* in either the intestine or body wall muscle can rescue the eRNAi body wall muscle silencing defect in *sid-1(qt95)* animals

The striking *sid-1(qt95)* phenotype is functional eRNAi limited to the intestine (Figure 3; Figure 7). Although this phenotype mimics a defect in dsRNA export from the intestine, multiple experimental approaches do not support a role for SID-1 in dsRNA export. *In vivo* and *in vitro* characterization of this allele thus far indicates that this phenotype is due to impaired import of dsRNA, which can be overcome by increasing the dose of dsRNA. However, it is not clear whether the *sid-1(qt95)* mutation specifically affects SID-1 function in the intestine, or if it primarily disrupts dsRNA import in distal tissues.

To address this question, we used the *sid-2* intestine-specific promoter and the *myo-3* body wall muscle (bwm)-specific promoter (Okkema, Harrison et al. 1993) to express wild-type *sid-1* selectively in proximal (intestine) and distal (muscle) tissues in *sid-1(qt95)* animals. We assayed both transgenic strains by eRNAi using bacteria expressing the muscle-specific *unc-22* dsRNA. If *sid-1(qt95)* animals have impaired activity only in the intestine, then providing WT *sid-1* under the control of the *sid-2* intestine-specific promoter should be sufficient for rescue of the mutant phenotype.Conversely, expression of WT *sid-1* under the control of the *myo-3* bwm-specific promoter should not rescue the *qt95* mutant phenotype. Unexpectedly, we found that both constructs fully rescued the *unc-22* eRNAi defect from almost no twitching animals in *sid-1(qt95)* mutants to nearly 100% twitching animals in the transgenic lines (Figure 9A). To ask whether the rescue was equivalent in the intestine and the muscle, we tested the tissue-specific transgenic lines on a dilution series of *unc-22* dsRNA expressing bacteria. This experiment revealed that the tissue-specific rescue lines are less sensitive than wild type at intermediate and low doses of dsRNA, but remain indistinguishable from each other at all dilutions (Figure 9B). Together, these results suggest a requirement for *sid-1* in both the proximal and distal tissues in the context of eRNAi. Alternatively, the transgene rescue constructs may be unintentionally and undetectably misexpressed.

**Figure 9.**
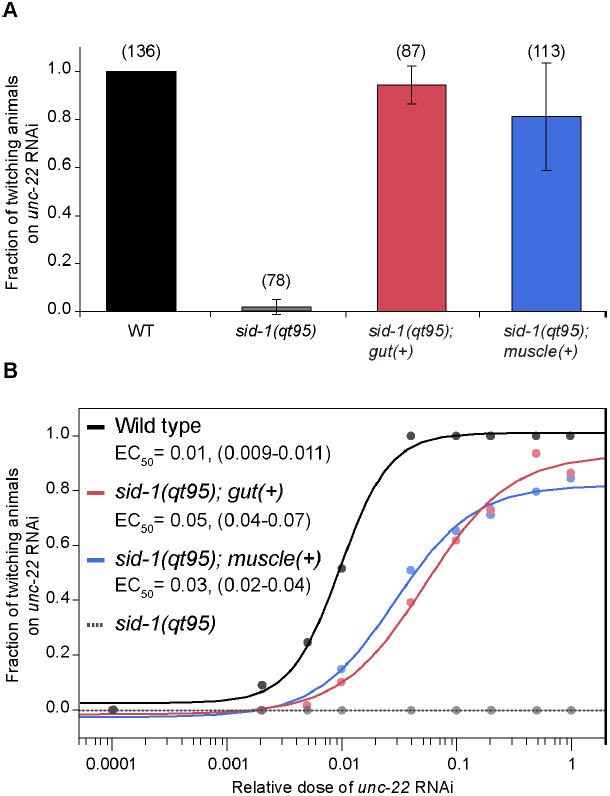
*sid-1(qt95)* animals have a weak dsRNA import defect that can be overcome by increased amounts of dsRNA. (A) RNAi sensitivity of *sid-1(qt95)* animals expressing SID-1(WT) under the control of a gut-specific promoter (Psid-2) or a muscle-specific promoter (Pmyo-3) compared to WT and sid-1(qt95) control animals. Number of animals scored (n) is shown for each condition. Error bars indicate standard deviation. (B) RNAi sensitivity in response to a dilution series of unc-22 RNAi The mean fraction of animals twitching and the apparent EC50 with 95% confidence interval in parentheses is shown. n > 40 animals for each strain at each dilution.

To verify that wild-type *sid-1* activity in muscle cells is sufficient to rescue eRNAi targeting muscle cells in *sid-1(qt95)* mutants under the native *sid-1* promoter we used genetic mosaic analysis. This experimental approach controls for misexpression because the transgene is either present or absent in the target tissue. Similar to the genetic mosaics described above, we co-injected *sid-1* wild-type genomic PCR product and a *sur-5::gfp* plasmid into *sid-1(qt95)* animals. We then identified by visual inspection, GFP positive animals that lacked GFP in the large, readily visible intestinal nuclei. These animals *are sid-1(qt95)* in the intestinal cells and *sid-1*(+) in other cells.These mosaic animals were placed on bacteria expressing *unc-22* dsRNA and scored two days later for the Unc-22 twitching phenotype. Sixteen of 21 animals twitched when placed in levamisole. Among the five animals that did not twitch, bwm cell GFP was not detected. Similar results (15/18 animals twitching) were obtained in *sid-1(qt9)* animals expressing the *sid-1* rescue construct outside of the intestine, confirming previous promoter expression studies that concluded that *sid-1* is not required in the intestine for transport of ingested dsRNA silencing signals to intestinal distal tissues (Jose, Smith et al. 2009). These results support the tissue-specific promoter results, which indicated that *sid-1(qt95)* disrupts *sid-1* activity in both proximal (intestine) and distal (muscle) cells.

## Discussion

Here we extend our genetic analysis of the novel dsRNA transport protein SID-1 (Feinberg and Hunter 2003, Shih, Fitzgerald et al. 2009, Shih and Hunter 2011). Our analysis of *sid-1* alleles shows that missense mutations that disrupt systemic RNAi almost exclusively alter conserved amino acids in transmembrane and extracellular domains. Within the N-terminal extracellular domain the mutations cluster near SID-1-specific domains that are missing in the closest homologs. These observations suggest that these domains support dsRNA transport in *C. elegans.* A disproportionate share of these ECD mutations cause a weak Sid phenotype. Because these mutations cause only a partial loss of function, their ability to transport dsRNA is not eliminated and their location in the ECD suggest they may disrupt interaction with either substrate (dsRNA) or other subunits. Indeed, the *in vitro* dsRNA binding affinity of purified SID-1 ECD is reduced 3-5 fold by SID-1(A173T) and SID-1(P199L) (Li et al., 2015). Furthermore, the Human SidT1 ECD forms tetramers in vivo and our functional analysis of SID-1 WT and SID-1(S536I) proteins is consistent with functional multimers (Pratt et al., 2012; Shih et al., 2009). This collection of missense alleles represents a strong starting point for additional biochemical studies investigating the mechanisms of substrate recognition and regulation of dsRNA channel activity.

A central challenge of this work was to understand the unique *sid-1(qt95)* eRNAi phenotype: *sid-1(qt95)* animals are fully sensitive to eRNAi in the gut, but cannot respond to eRNAi in any other tissue. Several lines of investigation support the hypothesis that this substitution disrupts export of silencing signals from the intestine, however the sum our data indicates that the apparent export defect reflects the cumulative effect of successive import defects. These data show that SID-1(D130N) expressed in *Drosophila* S2 cells is profoundly defective for dsRNA import (Figure 8).And consistent with an import defect, muscle-specific expression of wild-type *sid-1* in*sid-1(qt95)* animals restores effective eRNAi silencing in muscle cells (Figure 9).Although intestine-specific expression of wild-type *sid-1* in *sid-1(qt95)* animals also restores effective eRNAi silencing in muscle cells (Figure 9), our genetic mosaic analysis of *sid-1(qt9)(*null) mutants clearly shows that SID-1 is not required for export (Figure 6). A model consistent with all these data is that *sid-1*-dependent import within the intestine is important for subsequent efficient export (release) of dsRNA to intestine-distal tissues.

Our analysis of SID-1(D130N) activity in *Drosophila* S2 cells showed that increasing the dose (concentration) of dsRNA or exposure time to dsRNA could compensate for the reduced transport (Figure 8). Thus the observation that pseudocoelomic injected dsRNA overcomes the *sid-1(qt95)* defect (Figure 5) may be explained by the higher concentration of pseudocoelomic dsRNA following injection relative to eRNAi. Similarly, we found that *sid-1(qt95)* animals were more sensitive than wild type to a reduction in ingested dsRNA dose (Figure 9). Since the intestine is in direct contact with ingested dsRNA it may simply receive a higher ingested dsRNA dose relative to other tissues, which can receive only what the intestine releases (exports). In wild-type animals this difference in dose may be less apparent than in *sid-1(qt95)* mutants. Thus, the apparent tissue-specific deficiency observed in *sid-1(qt95)* animals may be explained by the cumulative effect of reduced dsRNA import, first in the intestine and then in the muscle.

The observation that increasing import activity in *sid-1(qt95)* animals by expressing wild-type SID-1 in either the intestine or the bwm cells is sufficient to partially restore eRNAi (Figure 9) indicates that *sid-1* functions in both tissues to support eRNAi. The function of SID-1 in bwm is to import silencing signals (Winston 2002). This is corroborated by the observations that target-tissue-specific SID-1 expression is sufficient for eRNAi in either *sid-1(qt9)* (Jose et al., 2009) or *sid-1(pk3321)* (Calixto et al., 2010) animals. These results were confirmed by our eRNAi analysis of genetic mosaic results reported here. What is the function of SID-1 in intestinal cells? Two models have been proposed for how SID-1 may function in the intestine to enable eRNAi (Whangbo and Hunter 2008). Common to both models is SID-2-dependent endocytosis of ingested dsRNA (McEwan, Weisman et al. 2012). In the first model SID-1 functions to transport ingested dsRNA directly into the intestinal cytoplasm, perhaps releasing dsRNA from endosomes. Once in the cytoplasm a SID-1-independent mechanism enables dsRNA export. This model explains how SID-1 expression in the intestine may boost dsRNA export to rescue the SID-1(D130N) impaired muscle import defect. In the second model endocytosed dsRNA is delivered to the pseudocoelom by vesicle fusion to the basal membrane (transcytosis). SID-1 then imports the pseudocoelomic dsRNA into distal tissues, including the intestine. This model is consistent with observed SID-1-independent transport of ingested dsRNA to *sid-1* expressing muscle cells (Jose et al., 2009; Calixto et al., 2010; Figure 9). SID-5 is required for this *sid-1*-independent pathway (Hinas, Wright et al. 2012). SID-5 is a small novel protein that co-localizes with late endosome/multivesicular body proteins and is implicated in export of eRNAi silencing signals (Hinas, Wright et al. 2012). Similar to our analysis of *sid-1(qt95)*, expressing wild-type *sid-5* in the intestine was sufficient to restore eRNAi silencing to body wall muscle cells. However, expressing wild-type *sid-5* in muscle cells was not sufficient to restore silencing, indicating that the *sid-5* defect is restricted to exporting cells. Furthermore, muscle-specific wild-type *sid-1* expression in*sid-1; sid-5* double mutants failed to restore silencing, indicating that *sid-5* is required for *sid-1*-independent transport of ingested dsRNA. Together with the results presented here, we can now conclude that *sid-1* and *sid-5* dependent pathways function together in the intestine to ensure efficient transport of ingested dsRNA to distal tissues for subsequent *sid-1*-dependent uptake.

## Materials and Methods

Transgenic animals for gut rescue of *sid-1* were generated using standard techniques and three independent lines were analyzed (Mello, Kramer et al. 1991). All images within a figure were adjusted in the same manner using ImageJ (NIH), and transferred into Illustrator (Adobe) for display. A list of primers used in this study is provided in Table S3.

### Strains

All strains were maintained using standard methods (Brenner 1974), and all assays were performed at 20°C. Bristol N2 was used as the wild type (Brenner 1974). See Table S4 for details of strains used in this study.

### Genetic mosaic analysis

*sid-1* genomic DNA was PCR amplified (primers C04F5.1 R1 and LGV W F, Table S3) from Bristol N2 worm extracts. *sid-1* gDNA and pTG96 (*sur-5::NLS-GFP*) were co-injected into *sid-1*(*qt9*) V mutants to create multiple transient lines. *sid-1* mosaic animals were directly identified by mosaic GFP expression patterns in the intestinal nuclei. For single cell injections, 200 ng/μL *gfp* dsRNA and 50 μM Alexa Fluor 555 conjugated to 10,000 MW Dextran (Molecular Probes) were mixed and injected into either GFP positive or negative cells of young adult animals.Injected worms were recovered to *E. coli* OP50 seeded NG plates for 1-2 hours before initial fluorescent and white-light imaging. To assess systemic silencing, animals were re-imaged 2 days (20°C) post-injection.

### Feeding RNAi assays

Feeding RNAi assays were adapted from standard procedures (Kamath and Ahringer 2003). Clones for each feeding RNAi vector (*act-5*/T25C8.2, *bli-1*/C09G5.6, *unc-22*/ZK617.1, *unc-45*/F30H5.1, *fkh-6*/B0286.5) were obtained from the Ahringer library (Kamath and Ahringer 2003). The *gfp* RNAi clone (pPD95.77) was a gift of A. Fire (Stanford University). Single colonies for each clone were inoculated in LB broth containing 100 μg/ml carbenicillin and 5 μg/ml tetracycline and grown overnight at 37°C. Overnight cultures for each clone were seeded onto 30 mm NG plates (Brenner 1974) containing 1mM isopropyl-β-D-thiogalactopyranoside (IPTG) and 100 μg/mL carbenicillin. After 24 hours at room temperature, RNAi plates were stored at 4°C for up to one month prior to use. L1-L2 staged larvae were placed on the RNAi plates and grown at 20°C for 2-3 days prior to scoring the RNAi phenotype. *act-5* RNAi animals are slow-growing and arrest at early larval stages (MacQueen et al., 2005). *bli-1* RNAi animals have fluid-filled blisters on the cuticle as adults (Brenner 1974). *unc-22* RNAi animals exhibit slow movement or paralysis with twitching body wall muscles. This twitching phenotype is enhanced by scoring the worms following submersion in 1 mM levamisole, as *unc-22* RNAi resistant animals become paralyzed and are motionless whereas *unc-22* RNAi sensitive animals are visibly twitching (Lewis et al., 1980). *unc-45* RNAi animals are paralyzed and sterile (Kachur, Audhya et al. 2008). *fkh-6* RNAi animals are sterile due to defects in the somatic gonad (Chang, Tilmann et al. 2004).

### dsRNA preparation and microinjection

Double-stranded *gfp* RNA was made by *in vitro* transcription using the AmpliScribe T7 High Yield Transcription Kit (Epicentre Technologies), and pPD95.77 as a template. Double-stranded *unc-22* RNA was made similarly using the Ahringer library clone as a template. Following extraction in Trizol (Life Technologies), all dsRNAs were annealed by heating to 90°C for 2 minutes and cooling one degree every eight seconds until reaching 25°C. All dsRNA injections used 100 ng/μl dsRNA in water with 1:200 diluted Alexa Fluor 555 conjugated to 10,000 MW Dextran (Molecular Probes) unless stated otherwise in the text. Late L4 and young adult stage hermaphrodite animals were microinjected using standard techniques and recovered at 20°C on OP50 seeded NG agar plates for 48 hours prior to scoring for the GFP silencing phenotype. *sid-1(-)* worms were always injected before *sid-1(+)* worms when using the same needle.

### Rescue of *sid-1* mutant by *cbn-sid-1*

To express *cbn*-*sid-1* in *sid-1(qt9)* animals, the *cbn-sid-1* promoter, gene (CBN00847) and 3’ UTR were amplified from *C. brenneri* (PB2801) gDNA in a single reaction with primers AW179 and AW180 (Table S3B). Purified PCR product (2 μl 25 ng/ul) was mixed with pHC183 (*myo-3*:dsRED) (1 μl 90ng/μl) was injected into HC196 to generate transgenic lines. Transgenic animals were identified by the presence of bwm RFP expression. Feeding assays with *unc-22* RNAi were performed and scored as described above.

### Gut and muscle rescue of *sid-1*

To express *sid-1*(+) cDNA in the intestine/gut of *sid-1(qt95)* animals, the *sid-2* promoter was amplified from gDNA with primers P23 and P37 and the *sid-1*(WT) cDNA and fused *unc-54* 3' UTR was amplified from pHC355 with primers P38 and P5 (Table S3D). A 1:1 mix of the two PCR products (0.037 mg/ml each) along with pHC183 (*myo-3*:dsRED) (0.037 mg/ml) was injected into HC726 to generate transgenic lines (Table S4). To express *sid-1(+)* cDNA in the body wall muscles (bwm) of *sid-1(qt95)* animals, we co-injected the plasmids pHC355 (*myo-2* promoter, *sid-1* cDNA, unc-54 3’UTR) (Jose, Smith et al. 2009) and pHC183 (*myo-3*:dsRED) into HC726 to generate temporary transgenic lines (Table S4. Transgenic animals were identified by the presence of bwm RFP expression. Feeding assays with *unc-22* RNAi were performed and scored as described above.

### Dilution series of *unc-22* and *gfp* dsRNA feeding

For dilution series, cultures of *E. coli* HT115(DE3) containing empty vector (pL4440) or *unc-22* RNAi were prepared in LB broth containing 100 μg/ml carbenicillin and 5 μg/ml tetracycline and grown overnight at 37°C. Optical density at 600 nm (OD600) values were determined by spectrophotometry and adjusted to 1.0. Bacteria were then mixed i n the following *unc-22* RNAi to pL4440 ratios: 0:1, 1:500, 1:200, 1:100, 1:25, 1:10, 1:5, 1:2, and 1:0. For each mix, 100 μLwas seeded onto 30 mm NG plates containing 1 mM IPTG and 100 μg/mL carbenicillin. Bacterial lawns were grown overnight at room temperature. Ten to twenty wild-type or transgenic *sid-1(qt95); gut::sid1(+)* or *sid-1(qt95); bwm::sid-1(+)* L4 stage animals were transferred to each type of RNAi plate and grown at 20°C. Animals were then scored for twitching in the presence of 1 mM levamisole 48 hours later. The fraction of twitching animals was calculated for each dilution of *unc-22* RNAi. Greater than 40 animals were scored for each condition. The half maximal effective concentration (EC_50_) was determined from fitting the data to a sigmoidal dose-response function using JMP Pro 11 (JMP Statistical Discovery Software from SAS). The EC_50_ was determined for each strain using the equation f=c+{(d-c)/(1+10^(-a(log_10_([P]-b))))}, where f is the fraction of animals twitching, [P] is the relative dilution of *unc-22* RNAi, b is the EC_50_, a is the Hill coefficient, d is the maximum fraction of animals twitching and c is the minimum fraction of animals twitching (10). Each experiment was performed at least in duplicate. The same procedure was repeated for the *gfp* RNAi dilution series using wild-type or *sid-1(qt95)* L4 stage animals carrying the *nrIs20 [sur-5::NLS-gfp]* transgene. Animals were scored for silencing of *gfp* expression in the intestinal nuclei at 48 hours. Comparison of EC50 values between strains was performed by analysis of means (ANOM) using JMP Pro 11.

### Microscopy of *C. elegans*

Images were acquired using HCImage software (Hamamatsu) and an Olympus SZX2 stereo microscope attached to a Lumen Dynamics X-Cite Fluorescence Illumination System for fluorescent detection assays. Animals were immobilized by chilling on NG agar plates to 4°C prior to photographing. After initial image capture, subsequent analysis of images was performed using ImageJ software (National Institutes of Health, USA). For display of the injected animals, two consecutive images were overlaid using ImageJ. The scale bar was measured using a hemocytometer. For Figure 6 bright field and fluorescent images were acquired using Openlab 3.1.7 (Improvision), a C4742-95 CCD (Hamamatsu) and a Zeiss Axiophot.

### Plasmid construction and Site-directed mutagenesis

Full length cbn-sid-1 was PCR amplified from mixed stage PB2801 cDNA (primers AW168 and AW169). Full length tag-130 (isoform A) was PCR amplified from mixed stage N2 cDNA (primers AW205 and AW206). FLAG^*^-pPacpl backbone-pAc5 was PCR amplified off pHC235 (primers AW170 and AW171). Fragments were gel extracted then assembled in Seamless Ligation Cloning Extract (SLiCE) (Zhang, Werling et al. 2014). Missense mutations were recreated by site-directed mutagenesis of a SID-1 cDNA expression plasmid, pHC235 using the Quikchange II XL kit (Stratagene) and primers listed in Table S3C.

### *Drosophila* S2 Cell Culture

S2 cells were cultured in Schneider's *Drosophila* medium (Invitrogen) supplemented with 10% fetal bovine serum and penicillin and streptomycin (Invitrogen). 48 hours prior to assays S2 cells were transiently transfected with 1 μg plasmid DNA using Effectene reagent (Qiagen).

### dsRNA synthesis for uptake assays in S2 cells

500 bp sense and antisense single T7 promoter templates were prepared as described (Feinberg and Hunter 2003). Radiolabeled 500 nt complementary single-stranded RNA were synthesized using ^32^P UTP (Perkin Elmer) and MAXIscript T7 *in vitro* transcription kit (Ambion/Invitrogen/Life Technologies). Unincorporated nucleotides were removed using Micro-Spin P-6 columns (Bio-Rad), percent incorporation determined by relative counts per minute and equal amounts of the single-strands were annealed as previously described (Feinberg and Hunter 2003). Unlabeled 500 nt ssRNAs were transcribed using Ampliscribe T7-Flash Transcription kit (Epicentre), extracted in Trizol (Life Technologies), precipitated with sodium acetate and ethanol, and redissolved in RNase-free 10mM Tris-HCl, pH 8.5. The OD260 of each strand was used to calculate the yield of each strand; equal amounts were mixed and annealed.

### dsRNA uptake assays in S2 cells

Standard assay: 4×10^6^ cells were pre-incubated at 4C for 0.5 hours prior to addition of dsRNA. Cells were incubated with dsRNA at 4C for 0.5 hrs. Cells were subsequently pelleted and resuspended in 0.1 ml of 0.25% Trypsin without EDTA (Gibco) for 8 minutes. Protease treatment was quenched upon addition of 1 ml ice-cold complete medium. Cells were pelleted and washed thoroughly three times with 1 ml ice-cold PBS, then lysed in 0.1% SDS. Samples were transferred to vials containing Ecoscint A (National Diagnostics) and scintillation was measured with a Beckman LS 6000 IC. Modified assays were performed as above unless otherwise stated: (a) Extended time assay: cells were incubated with ^32^P-dsRNA at 4C for 25 hrs; (b) Increased dsRNA assay: cells were co-incubated with 490 ng/ml unlabeled dsRNA and10 ng/ml ^32^P-dsRNA at 4C for 0.5 hrs; (c) Kinetic, short and intermediate assays: cells were maintained at room temperature and incubated with dsRNA for 1 to 50 minutes (as specified in figure legend).

### Immunofluorescence of *Drosophila* S2 cells

Transfected S2 cells were allowed to adhere to uncoated 35mm glass bottom dishes for 45 minutes in complete medium, fixed in 10% Formaldehyde in PBS for 20 minutes, rehydrated and permeabilized by 3 washes of PBS-0.1% Triton X-100. Samples were blocked with 5% normal goat serum in PBS-T for 30 minutes at room temperature. Monoclonal ANTI-FLAG M2 antibody produced in mouse (Sigma) was used as the primary antibody followed by Goat anti-Mouse IgG, Alexa Fluor 647 (Invitrogen) as the secondary. DNA was labeled with VECTASHIELD mounting medium with DAPI (Vector Labs). Fluorescent images were collected on an LSM 700 Inverted axio Observer (Zeiss) at the Harvard Center for Biological Imaging.Immunofluorescence of S2 cells expressing the various SID-1 homologs revealed that localization and expression of the proteins was variable from cell to cell, but indistinguishable across transfections (Figure S2).

### Sequencing of *sid-1* alleles

To identify the molecular lesion in *sid-1* mutants, the *sid-1* gene was amplified from mutant genomic DNA obtained from lysis of adult hermaphrodite worms. Mutant hermaphrodites (5-6 adults) were picked into separate PCR tubes, each containing 5 μL worm lysis buffer (Qiagen Buffer EB, 150 μg/ml proteinase K). Worms were lysed by freezing in liquid nitrogen followed by incubation at 65°C for 1 hour and incubation at 95°C for 15 minutes (for inactivation of proteinase). The lysate DNA templates were then immediately combined with 45 μL of a PCR master mix containing: 1.75 μL 10 mM dNTPs, 1.5 μL 10μM LGV W Forward (forward primer),1.5 μL 10μM C04F5.1 Reverse 1 (reverse primer), 0.75 μL Expand Long Template Enzyme Mix (Roche), 5 μL 10x Expand Long Template Buffer 2 and 34.5 μL water. PCR reactions were done using the cycling conditions: 92°C for 2 minutes, 10 cycles of (92°C for 10 seconds, 55°C for 30 seconds, 68°C for 8 minutes), 22 cycles of (92°C for 10 seconds, 55°C for 30 seconds, 68°C for 8 minutes + 20 seconds/cycle), 68°C for 7 minutes. After amplification, the PCR products were run on a 0.8% agarose gel and the 7.8 kb *sid-1* genomic DNA band was extracted and purified using the QIAquick Gel Extraction Kit. The purified product was eluted in 50 μL Buffer EB and sent for sequencing at the Dana-Farber/Harvard Cancer Center DNA Resource Core. Sequences were compared to the wild-type *sid-1* sequence reported by Winston et al. (2002) (GenBank AF478687). See Table S3A for details of primers used for sequencing. Two independent amplifications were sequenced for each allele.

### SID-1 phylogeny, percent identity matrix and sequence alignment

Phylogenetic neighbor-joining tree, percent identity matrix and clustalW-formatted alignment of *C. elegans* SID-1 and eight *Caenorhabditis* homologs were generated using the MUltiple Sequence Comparison by Log-Expectation (MUSCLE) 3.8 multiple sequence alignment program for proteins (Edgar, 2004). Protein sequences were obtained from WormBase.org v258, WormBase ID numbers are listed below. The *C. brenneri* CHUP-1 sequence was edited (^*^) to remove a false exon present in the protein sequence [SRASSVSIMENSQSSRLNGGHR]. No full length CHUP-1 homolog has yet been identified in *C. japonica*; a partial sequence (56% length) was excluded from the alignment.

CEL-SID-1 WP:CE30331

CBR-SID-1 BP:CBP22114

CBN-SID-1 CN:CN02065

CRE-SID-1 RP:RP37195

CJP-SID-1 JA:JA60059

CEL-CHUP-1 WP:CE26397

CBN-CHUP-1^*^ CN:CN03240^*^

CBR-CHUP-1 BP:CBP03753

CRE-CHUP-1 RP:RP29620

## Acknowledgements

We thank Olga Minkina, Phil Shiu, and Javier Apfeld for comments on this manuscript, the Hunter lab for numerous discussions, the Harvard Center for Biological Imaging for infrastructure and support, and Wormbase. Some strains were provided by the Caenorhabditis Genetics Center, which is funded by NIH Office of Research Infrastructure Programs (P40 OD010440). This work was supported by NIH grant GM089795.

---

**Figure S1.**
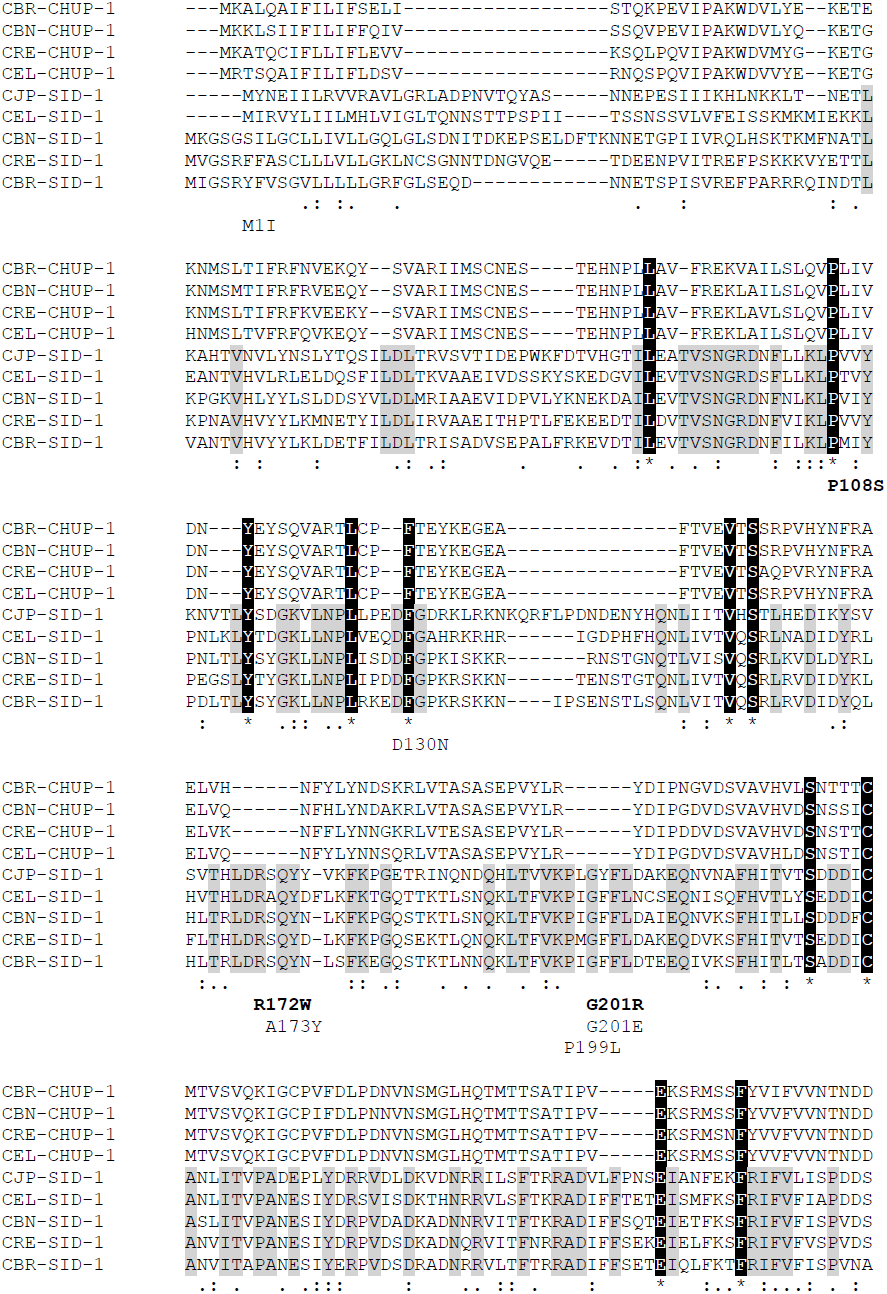
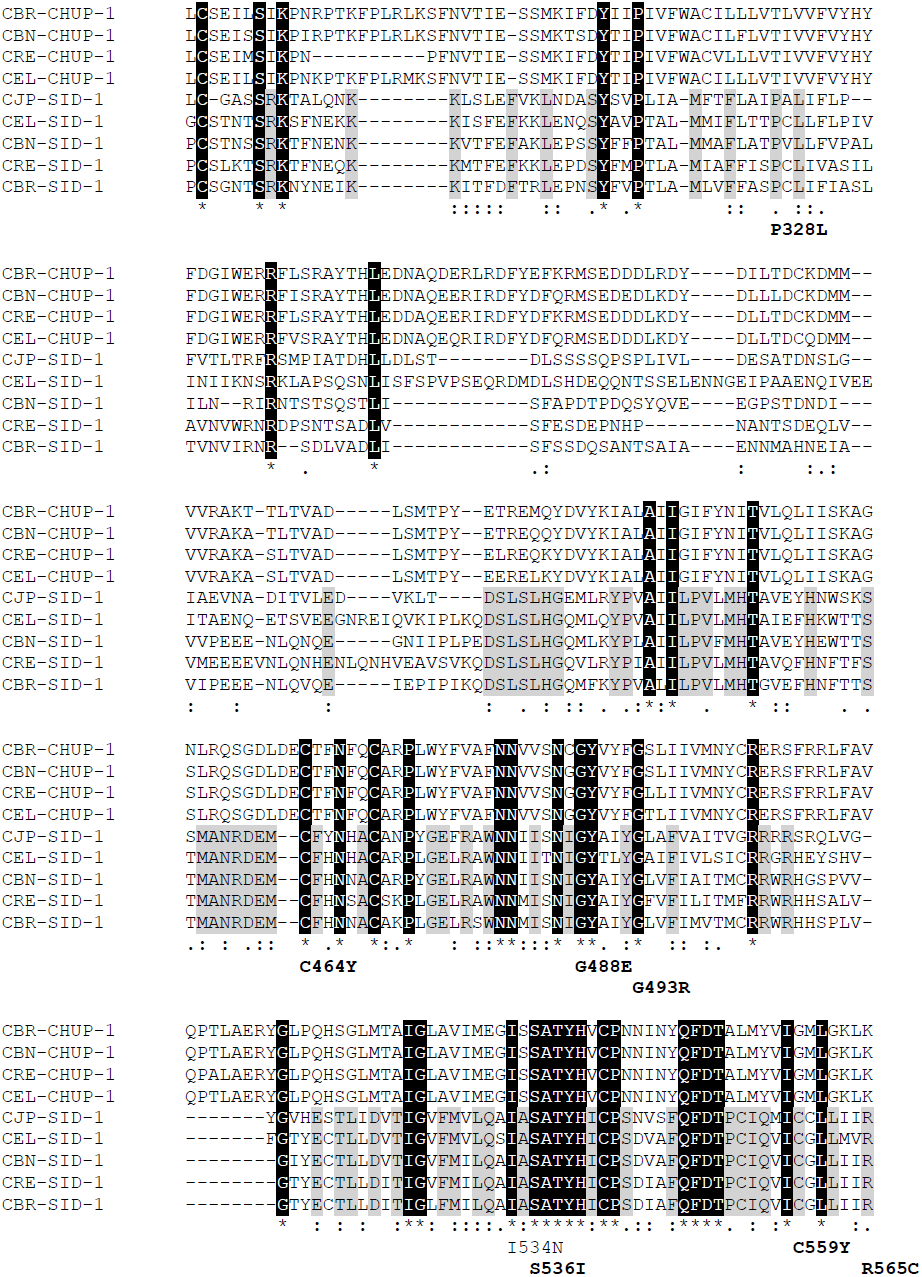
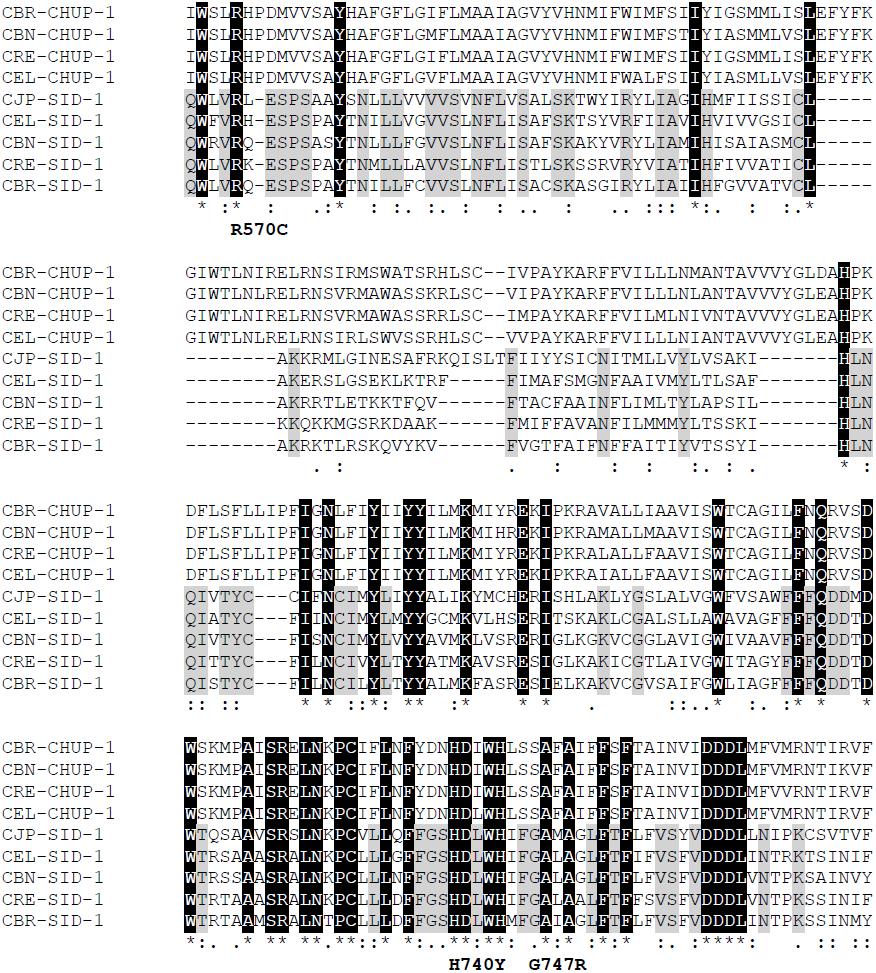
A multiple sequence alignment of full-length SID-1 homologs. Available full-length SID-1 and CHUP-1 homologs from *C. elegans, C. brenneri, C. briggsae, C. remanei,* and *C. japonica* were aligned by MUSCLE v3.8 with default parameters edgar (Edgar 2004). Residues identically conserved in all nine aligned sequences (predicted SID-1 orthologs and paralogs) are highlighted in black; residues identically conserved only in the five putative SID-1 orthologs are highlighted in grey. The position and amino acid change of the recovered weak alleles (regular weight) and strong alleles (bold weight) is indicated below the substituted residue.

**Figure S2.**
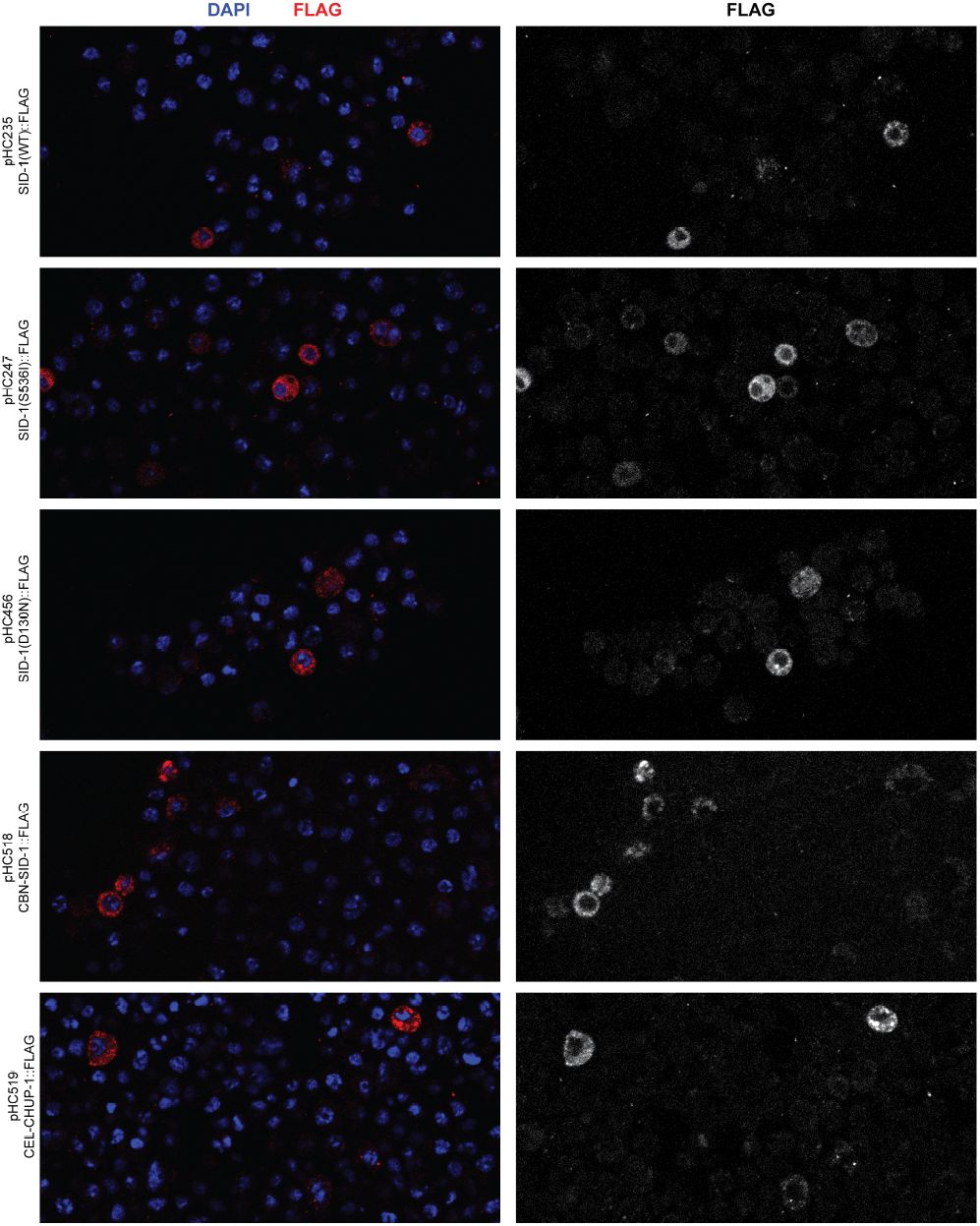
Anti-FLAG immunofluorescence of S2 cells expressing the indicated SID-1 constructs. Each image presented side by side as composite and single channel: in composite (left) FLAG is displayed in the red, DNA displayed in blue; FLAG only (right). Fluorescent images were collected on an LSM 700 Inverted axio Observer (Zeiss) at the Harvard Center for Biological Imaging. All images are adjusted identically.

**Table S1.**
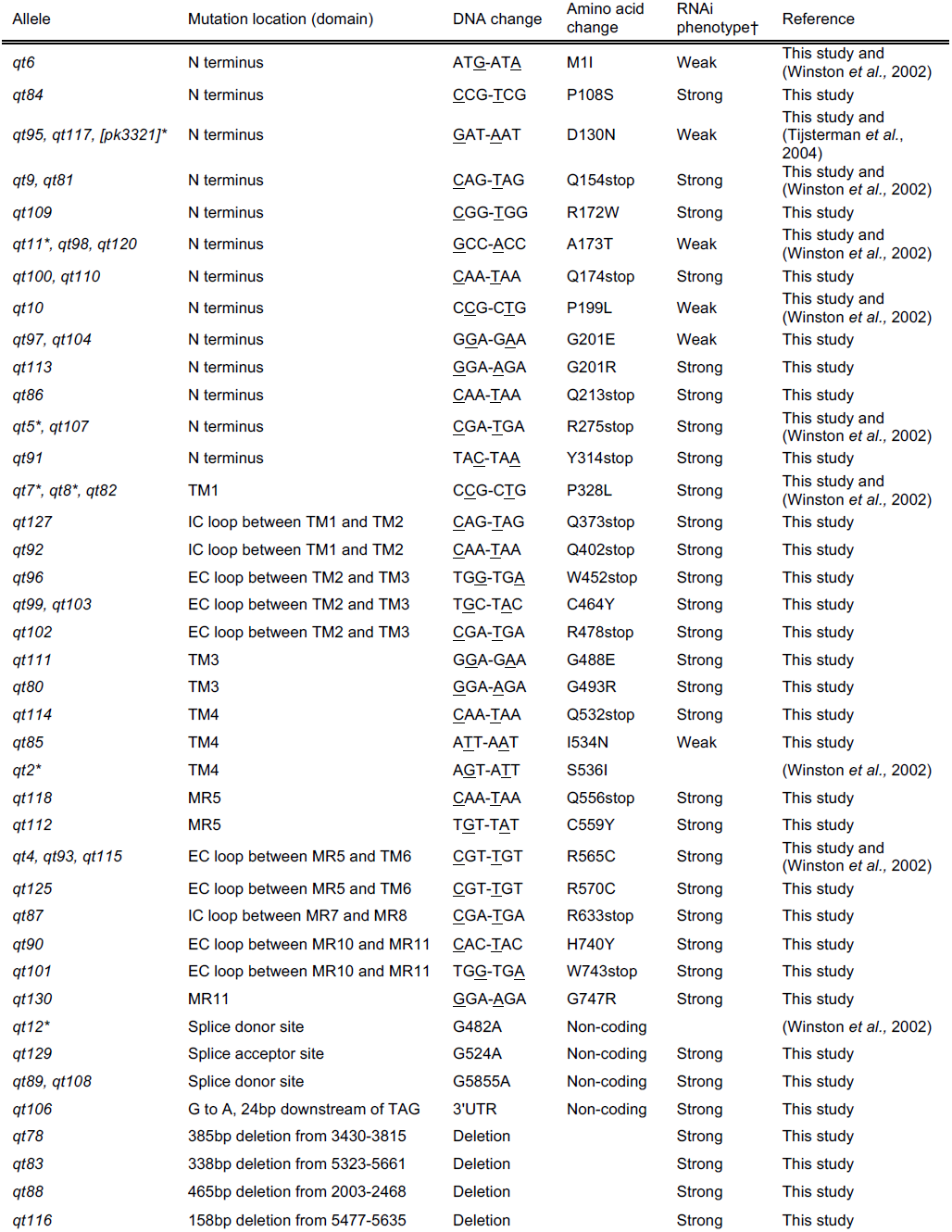
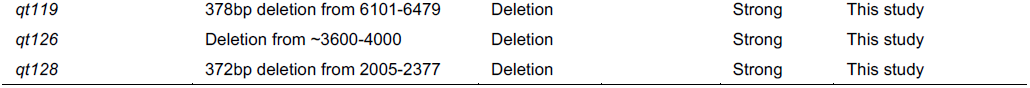
Sequence analysis of *sid-1* alleles. TM, transmembrane region; MR, membrane region; EC, extracellular; IC, intracellular. Asterisk indicates alleles that were not tested on multiple foods in this study. †The strength of the Sid phenotype was defined by the response to a panel of environmental RNAi assays: strong, fewer than 10% of animals sensitive to each tested trigger; weak, all other response profiles.

**Table S2.** Strong *sid-1* alleles on tissue-specific environmental RNAi. N2, wild type strain. HC57, background strain for recovery of *sid-1* mutants. Bli, blistered cuticle; Unc, uncoordinated (*unc-45* RNAi animals are paralyzed, *unc-22* RNAi animals are twitching). Fertility was scored qualitatively: sterile (no progeny), fertile (many progeny), decreased (few progeny). (n), number of animals scored. ND, not determined.

| Mutation | Strain or Allele | *act-5* RNAi<br>gut |  | *bli-1* RNAi<br>hypodermis |  | *unc-22* RNAi<br>muscle |  | *unc-45* RNAi<br>muscle, germline |  |  | *fkh-6* RNAi<br>somatic gonad |  |
| --- | --- | --- | --- | --- | --- | --- | --- | --- | --- | --- | --- | --- |
|  |  | % Arr | (n) | % Bli | (n) | % Unc | (n) | % Unc | Fertility | (n) | Fertility | (n) |
| CONTROLS |  |  |  |  |  |  |  |  |  |  |  |  |
|  | N2 | 100 | (111) | 100 | (115) | 100 | (173) | 100 | Sterile | (237) | Sterile | (173) |
|  | HC57 | 100 | (64) | 100 | (34) | 100 | (43) | 100 | Sterile | (48) | Sterile | (12) |
| NONSENSE AND NON-CODING, STRONG |  |  |  |  |  |  |  |  |  |  |  |  |
| Q154stop | *qt9* | 0 | (98) | 0 | (37) | 0 | (60) | 0 | Fertile | (24) | Fertile | (35) |
| Q154stop | *qt81* | 0 | (57) | 0 | (33) | 0 | (58) | 0 | Fertile | (68) | Fertile | (24) |
| Q174stop | *qt100* | 0 | (18) | 6 | (32) | 0 | (35) | 0 | Fertile | (24) | Fertile | (29) |
| Q174stop | *qt110* | 0 | (24) | 0 | (19) | 0 | (81) | 0 | Fertile | (74) | Fertile | (50) |
| Q213stop | *qt86* | 0 | (50) | ND |  | 0 | (31) | 0 | Fertile | (62) | Fertile | (57) |
| R275stop | *qt107* | 0 | (18) | 0 | (20) | 0 | (48) | 0 | Fertile | (42) | Fertile | (47) |
| Y314stop | *qt91* | 0 | (25) | ND |  | 0 | (51) | 0 | Fertile | (33) | Fertile | (38) |
| Q373stop | *qt127* | 0 | (23) | 0 | (17) | 0 | (19) | 0 | Fertile | (14) | Fertile | (17) |
| Q402stop | *qt92* | 0 | (16) | 6 | (33) | 0 | (17) | 0 | Fertile | (31) | Fertile | (33) |
| R478stop | *qt102* | 0 | (20) | 0 | (19) | 0 | (18) | 0 | Fertile | (22) | Fertile | (18) |
| W452stop | *qt96* | 0 | (35) | 0 | (34) | 6 | (35) | 0 | Fertile | (30) | Fertile | (35) |
| Q532stop | *qt114* | 0 | (27) | 0 | (22) | 0 | (22) | 0 | Fertile | (24) | Fertile | (18) |
| Q556stop | *qt118* | 0 | (23) | 0 | (9) | 0 | (15) | 0 | Fertile | (14) | Fertile | (25) |
| R633stop | *qt87* | 0 | (43) | ND |  | 0 | (44) | 0 | Fertile | (41) | Fertile | (35) |
| W743stop | *qt101* | 0 | (20) | 0 | (33) | 0 | (30) | 0 | Fertile | (27) | Fertile | (41) |
| Δ2003-2468 | *qt88* | 0 | (65) | ND |  | 0 | (58) | 0 | Fertile | (50) | Fertile | (41) |
| Δ2005-2377 | *qt128* | 0 | (14) | 0 | (18) | 0 | (23) | 0 | Fertile | (20) | Fertile | (16) |
| Δ3430-3815 | *qt78* | 0 | (30) | 0 | (10) | 0 | (16) | 4 | Fertile | (23) | Fertile | (11) |
| ~Δ3600-4000 | *qt126* | 0 | (21) | 0 | (17) | 0 | (21) | 0 | Fertile | (14) | Fertile | (11) |
| Δ5323-5661 | *qt83* | 0 | (57) | 0 | (53) | 0 | (67) | 0 | Fertile | (63) | Fertile | (32) |
| Δ5477-5635 | *qt116* | 0 | (25) | 0 | (26) | 0 | (18) | 0 | Fertile | (14) | Fertile | (21) |
| Δ6101-6479 | *qt119* | 0 | (22) | 0 | (25) | 0 | (44) | 0 | Fertile | (32) | Fertile | (29) |
| SAS G524A | *qt129* | 0 | (28) | 0 | (33) | 0 | (27) | 0 | Fertile | (30) | Fertile | (21) |
| SDS G5855A | *qt89* | 0 | (33) | ND |  | 0 | (43) | 0 | Fertile | (34) | Fertile | (35) |
| SDS G5855A | *qt108* | 0 | (18) | 0 | (12) | 0 | (56) | 0 | Fertile | (45) | Fertile | (28) |
| 3'UTR G6649A | *qt106* | 0 | (12) | 0 | (14) | 0 | (39) | 0 | Fertile | (40) | Fertile | (13) |
| MISSENSE, STRONG |  |  |  |  |  |  |  |  |  |  |  |  |
| P108S | *qt84* | 0 | (75) | 0 | (31) | 0 | (66) | 0 | Fertile | (60) | Fertile | (32) |
| R172W | *qt109* | 0 | (24) | 0 | (18) | 0 | (34) | 0 | Fertile | (56) | Fertile | (35) |
| G201R | *qt113* | 0 | (30) | 0 | (20) | 0 | (33) | 0 | Fertile | (14) | Fertile | (18) |
| P328L | *qt82* | 0 | (21) | 0 | (27) | 0 | (33) | 0 | Fertile | (28) | Fertile | (9) |
| C464Y | *qt99* | 0 | (20) | 0 | (36) | 0 | (36) | 0 | Fertile | (42) | Fertile | (33) |
| C464Y | *qt103* | 0 | (20) | 0 | (41) | 3 | (35) | 0 | Fertile | (38) | Fertile | (24) |
| G488E | *qt111* | 0 | (9) | 0 | (20) | 0 | (44) | 0 | Fertile | (51) | Fertile | (47) |
| G493R | *qt80* | 0 | (48) | 0 | (51) | 0 | (86) | 0 | Fertile | (52) | Fertile | (45) |
| C559Y | *qt112* | 0 | (28) | 0 | (23) | 0 | (50) | 0 | Fertile | (53) | Fertile | (48) |
| R565C | *qt93* | 0 | (28) | 0 | (20) | 0 | (24) | 0 | Fertile | (23) | Fertile | (16) |
| R565C | *qt115* | 0 | (36) | 0 | (37) | 0 | (24) | 0 | Fertile | (27) | Fertile | (31) |
| R570C | *qt125* | 0 | (38) | 0 | (15) | 0 | (28) | 0 | Fertile | (19) | Fertile | (16) |
| H740Y | *qt90* | 0 | (46) | ND |  | 0 | (45) | 0 | Fertile | (41) | Fertile | (46) |
| G747R | *qt130* | 0 | (18) | 0 | (29) | 0 | (33) | 0 | Fertile | (38) | Fertile | (24) |

**Table S3.**
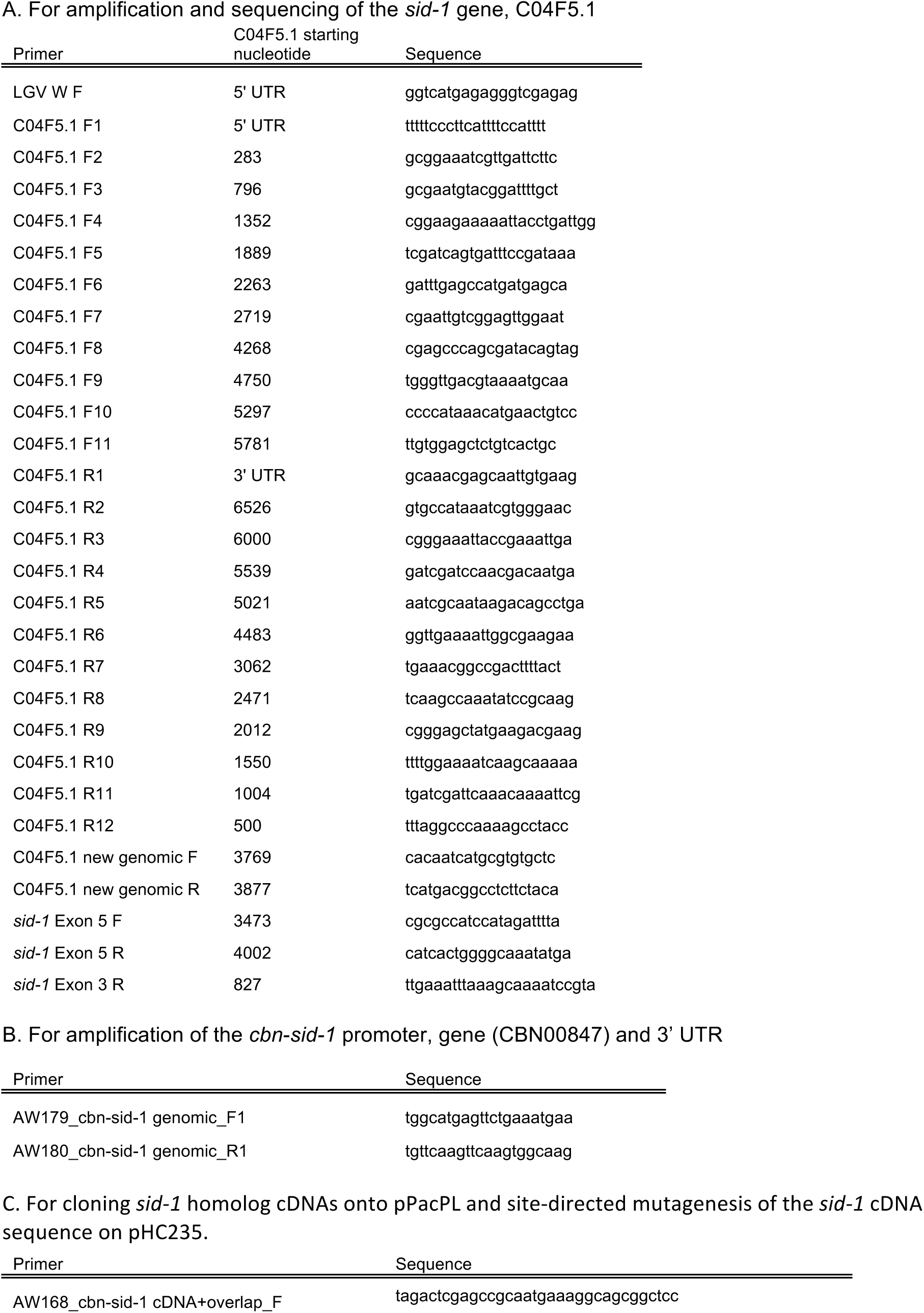
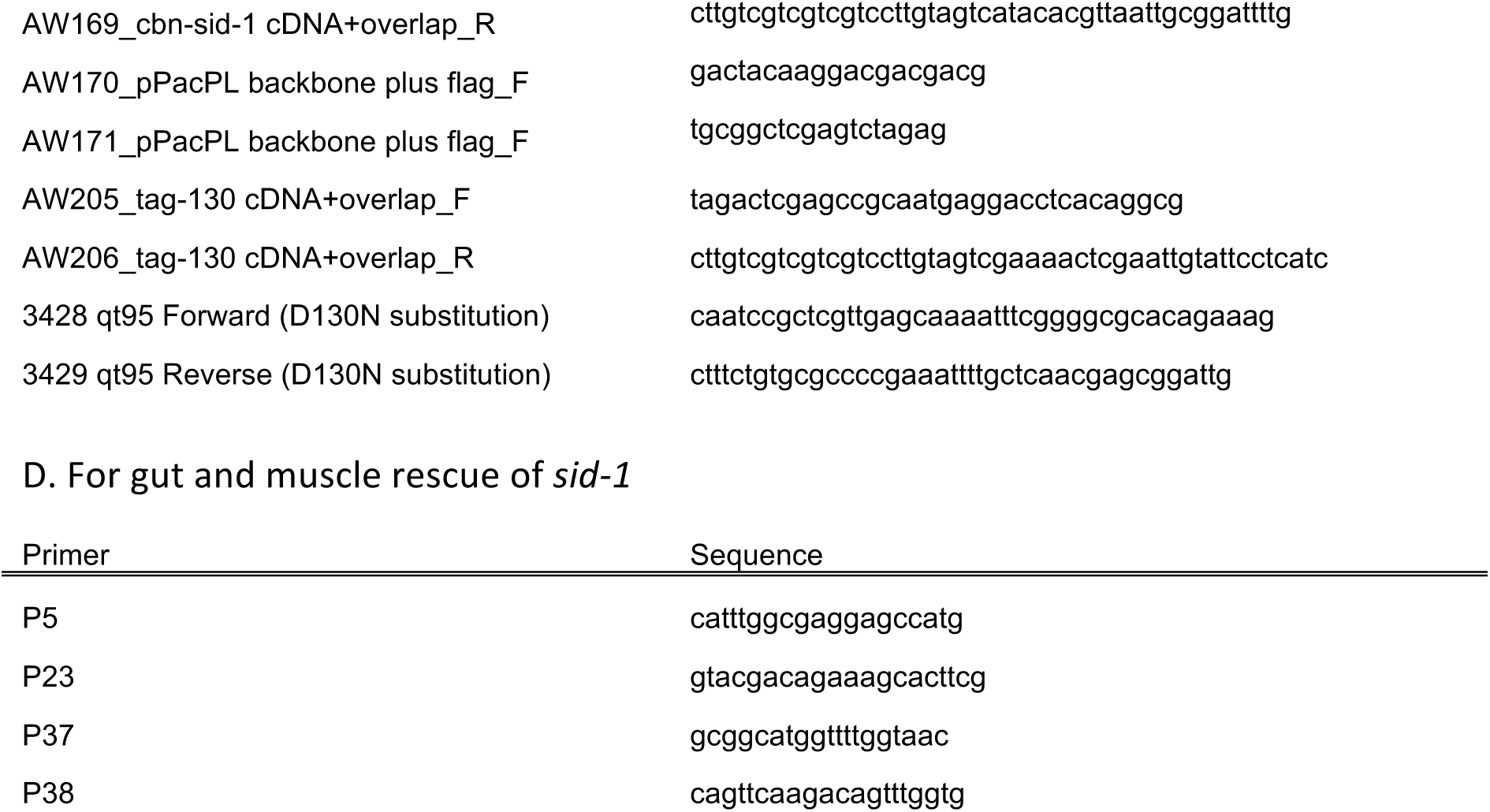
Sequences of primers used in this study.

**Table S4.** Strains used in this study

| Strain | Genotype |
| --- | --- |
| N2 Bristol | wild type |
| HC195 | *nrls20* [*sur-5::NLS-gfp*] IV |
| HC196 | *sid-1(qt9)* V |
| HC393 | *sid-1(qt9)* V; *qtEx89* [*sur-5::NLS-gfp* + *sid-1(+)*] |
| HC566 | *nrls20* [*sur-5::NLS-gfp*] IV; *sid-1(qt9)* V |
| HC757 | *nrls20* [*sur-5::NLS-gfp*] IV; *sid-1(qt95)* V |
| HC57 | *ccls4251*[*myo-3::NLS-gfp*, *myo-3::MITO-gfp*] I; *qtIs3* [*myo-2::GFP dsRNA*] III; *mls11* [*myo-2::GFP*] IV |
| HC303 | *sid-1(qt6)* V |
| HC94 | *sid-1(qt10)* V |
| HC653 | *sid-1(qt78)* V |
| HC655 | *sid-1(qt80)* V |
| HC656 | *sid-1(qt81)* V |
| HC657 | *sid-1(qt82)* V |
| HC658 | *sid-1(qt83)* V |
| HC659 | *sid-1(qt84)* V |
| HC660 | *sid-1(qt85)* V |
| HC661 | *sid-1(qt86)* V |
| HC662 | *sid-1(qt87)* V |
| HC663 | *sid-1(qt88)* V |
| HC664 | *sid-1(qt89)* V |
| HC665 | *sid-1(qt90)* V |
| HC666 | *sid-1(qt91)* V |
| HC667 | *sid-1(qt92)* V |
| HC668 | *sid-1(qt93)* V |
| HC670 | *sid-1(qt95)* V |
| HC671 | *sid-1(qt96)* V |
| HC672 | *sid-1(qt97)* V |
| HC673 | *sid-1(qt98)* V |
| HC674 | *sid-1(qt99)* V |
| HC675 | *sid-1(qt100)* V |
| HC676 | *sid-1(qt101)* V |
| HC677 | *sid-1(qt102)* V |
| HC678 | *sid-1(qt103)* V |
| HC679 | *sid-1(qt104)* V |
| HC681 | *sid-1(qt106)* V |
| HC682 | *sid-1(qt107)* V |
| HC683 | *sid-1(qt108)* V |
| HC684 | *sid-1(qt109)* V |
| HC685 | *sid-1(qt110)* V |
| HC686 | *sid-1(qt111)* V |
| HC687 | *sid-1(qt112)* V |
| HC688 | *sid-1(qt113)* V |
| HC689 | *sid-1(qt114) V* |
| HC690 | *sid-1(qt115) V* |
| HC691 | *sid-1(qt116) V* |
| HC692 | *sid-1(qt117) V* |
| HC693 | *sid-1(qt118) V* |
| HC694 | *sid-1(qt119) V* |
| HC709 | *sid-1(qt120) V* |
| HC714 | *sid-1(qt125) V* |
| HC715 | *sid-1(qt126) V* |
| HC716 | *sid-1(qt127) V* |
| HC717 | *sid-1(qt128) V* |
| HC718 | *sid-1(qt129) V* |
| HC719 | *sid-1(qt130) V* |
| HC935 | *sid-1(qt95) V; qtEx157[sid-2::sid-1 cDNA, myo-3::dsRed]* |
| HC936 | *sid-1(qt95) V; qtEx157[sid-2::sid-1 cDNA, myo-3::dsRed]* |
| HC937 | *sid-1(qt95) V; qtEx158[myo-3::sid-1 cDNA, myo-3::dsRed]* |
| HC938 | *sid-1(qt95) V; qtEx158[myo-3::sid-1 cDNA, myo-3::dsRed]* |
| HC1034 | *sid-1(qt9) V; qtEx195[cbn-sid-1 promoter::cbn-sid-1::3'UTR; pHC183 myo-3::dsRed]* |
| PB2801 | *Caenorhabditis brenneri*, outcrossed x20 |

Strains HC94 and HC653 through HC719 are in the HC57 strain background.

